# Human spermatogenesis leads to a reduced nuclear pore structure and function

**DOI:** 10.1101/2024.10.30.620797

**Authors:** Ália dos Santos, Oliver Knowles, Tom Dendooven, Thomas Hale, Victoria L Hale, Alister Burt, Piotr Kolata, Giuseppe Cannone, Dom Bellini, David Barford, Matteo Allegretti

## Abstract

Nuclear pore complexes (NPCs) are nuclear gateways which regulate the transit of molecules larger than 40kDa through a Ran-dependent transport^1^. The somatic human NPC scaffold consists of three stacked rings: the cytoplasmic (CR), the nucleoplasmic (NR), and the inner ring (IR), which define a central channel approximately 55nm wide^2^.

Although many studies have investigated human NPC architecture^3^, it remains largely unknown how the NPC accommodates the different functions of non-somatic cells.

Here, we reveal the *in-cell* architecture of the human sperm NPC. We show that it exhibits a central channel less than 40nm-wide, outlined exclusively by the IR. This structural alteration is accompanied by a six-fold reduction in nuclear diffusion rate and mis-localization of Ran-dependent transport components. Additionally, we identify a network of septin filaments interconnecting NPCs within the inter-membrane space of the nuclear envelope (NE), suggesting a potential mechanical role in channel constriction. Furthermore, our human tissue imaging data indicate meiosis as a pivotal differentiation stage driving architectural changes to the NPC.

Given the critical role of the IR for successful spermatogenesis^4,5^, our work offers important insight into this fundamental biological process. Our study also demonstrates the power of an integrative approach by combining electron-cryo- tomography, super-resolution-light-microscopy and biochemical analysis, to elucidate macromolecular structure-function relations in human physiological contexts.

## Introduction

In eukaryotic cells, the nucleus stores and protects the genetic information by enclosing it within a double lipid membrane, the nuclear envelope (NE). At the NE, large 110MDa protein complexes, known as nuclear pore complexes (NPCs), stabilise holes and allow regulated bidirectional transport and material exchange between the nucleus and the cytoplasm. NPCs are therefore crucial in the regulation of gene expression^6–8^.

The structure of NPCs consists of three stacked multi-subunit rings surrounding a central channel of approximately 55nm diameter: two outer rings - the cytoplasmic and nuclear rings (CR and NR, respectively) - and the inner ring (IR), which sits at the point of fusion between the two lipid membranes. An additional ring, the luminal ring (LR), lies within the intermembrane space, around the IR^3,9^ (Extended Data Fig. 1a). Each of these rings consists of multiple proteins, known as nucleoporins (NUPs), organised as supramolecular assemblies. In humans, scaffold NUPs form the IR and two Y-complex rings that lie at the interface with both the CR and NR^10^. Additional proteins anchor to the Y-complex to form either the cytoplasmic filaments at the CR side of the NPC^2,11–13^ at later stages of assembly^14,15^, or to form the nuclear basket that extends from the NR into the nucleoplasm^16^ (Extended Data Fig. 1a). The nuclear basket is critical for mRNA export^17^ and nuclear import^18^.

Several CR, NR and IR subunits contain phenylalanine-glycine (FG) repeats that extend towards the middle of the central channel. Molecules below 40kDa in size can passively diffuse through the NPC, whilst transport of larger proteins is highly regulated^1,19^. This is achieved through interactions of FG repeats at the central channel with transport receptors and the small GTPase Ran^1^. Importantly, Ran determines transport directionality by creating a Ran gradient that is dependent on its nucleotide status – enrichment of Ran-GTP in the nucleus and Ran-GDP in the cytoplasm^1^.

At the CR side of the NPC, RanBP2, also known as NUP358, stabilises two Y-complex-ring assemblies^20^ and binds to RanGAP, the Ran-activating protein, which ensures enrichment of Ran-GDP in the cytoplasm. RanBP2 is therefore critical in the regulation of nucleocytoplasmic trafficking^21^. Additionally, RanBP2 is a central component of NPCs present at stacked endoplasmic reticulum (ER) sheets, known as annulate lamellae^22–24^. These structures have been proposed to act as NPC storage pools during postmitotic NE assembly^25^ and animal development^22,23^.

NPCs are dynamic complexes capable of dilating and contracting to adapt to their mechanical environments^2,26–31^. This has been shown under several physiological conditions^26,31–33^ and during differentiation^34,35^. NPCs extracted from cells^20^, at annulate lamellae^24^, or in energy-depleted cells^31^ display the highest reported levels of constriction, with a central channel diameter smaller than 50nm. NPC constriction has been also associated to slower passive diffusion of small molecules into the nucleus^29^. Among mechanical forces acting on the NE, membrane tension, modulated by cytoskeletal forces, for example, is considered to be a major contributor to changes in NPC dynamics^31,32,36^. One prominent example of this phenomenon is spermiogenesis, the post-meiotic differentiation of male germ cells in testes. During this process, the nucleus of spermatids (post-meiotic cells) undergoes radical changes that culminate in the formation of mature spermatozoa capable of delivering the paternal DNA to the oocyte during fertilization^37,38^. These changes include DNA hyper- condensation through the replacement of histones with protamines, and elongation of the nucleus into a drop-shaped structure^37,38^. This is accompanied by time-sensitive regulation of protein expression, transcription shutdown and downregulation of nuclear transport^39,40^. Importantly, NPCs are also redistributed to the caudal region of the sperm head into redundant NE regions^41^. Previous work has suggested that several NUPs are critical during spermiogenesis^42,43^, in particular NUPs of the IR^4,5^. However, it is unknown whether NPC architecture is altered in germ cells or if structural changes would affect nuclear trafficking. This knowledge is particularly relevant to the study of early embryo development, as the whole paternal nucleus is delivered intact to the oocyte during fertilisation^44^ .

Here, we investigate the architecture and function of human sperm NPCs using an *in situ* integrative structural biology approach. We use a combination of electron-cryo- tomography (cryo-ET), super-resolution light microscopy, immunohistochemistry of human primary tissue and biochemical approaches, to reveal a highly constricted and reduced NPC scaffold with altered nuclear transport in human sperm.

## Results

### Clustering of NPCs in human sperm cells

In somatic cells, NPCs are distributed ubiquitously across the NE (Extended Data Fig. 1b). To investigate the localisation of NPCs in human sperm cells, we analysed samples from 25 healthy human donors. Using the MAb414 antibody as a marker for NPCs, we imaged human sperm cells with Structured Illumination Microscopy (SIM). Similarly to what was previously shown^41^, we confirmed that NPCs are exclusively localised at the rear of the nucleus (Fig. 1a). 3D SIM reconstruction revealed that NPCs form a single pocket surrounding the rear of the sperm head and flanking the tail (Fig. 1b and Extended Data Fig. 1c).

**Fig. 1.**
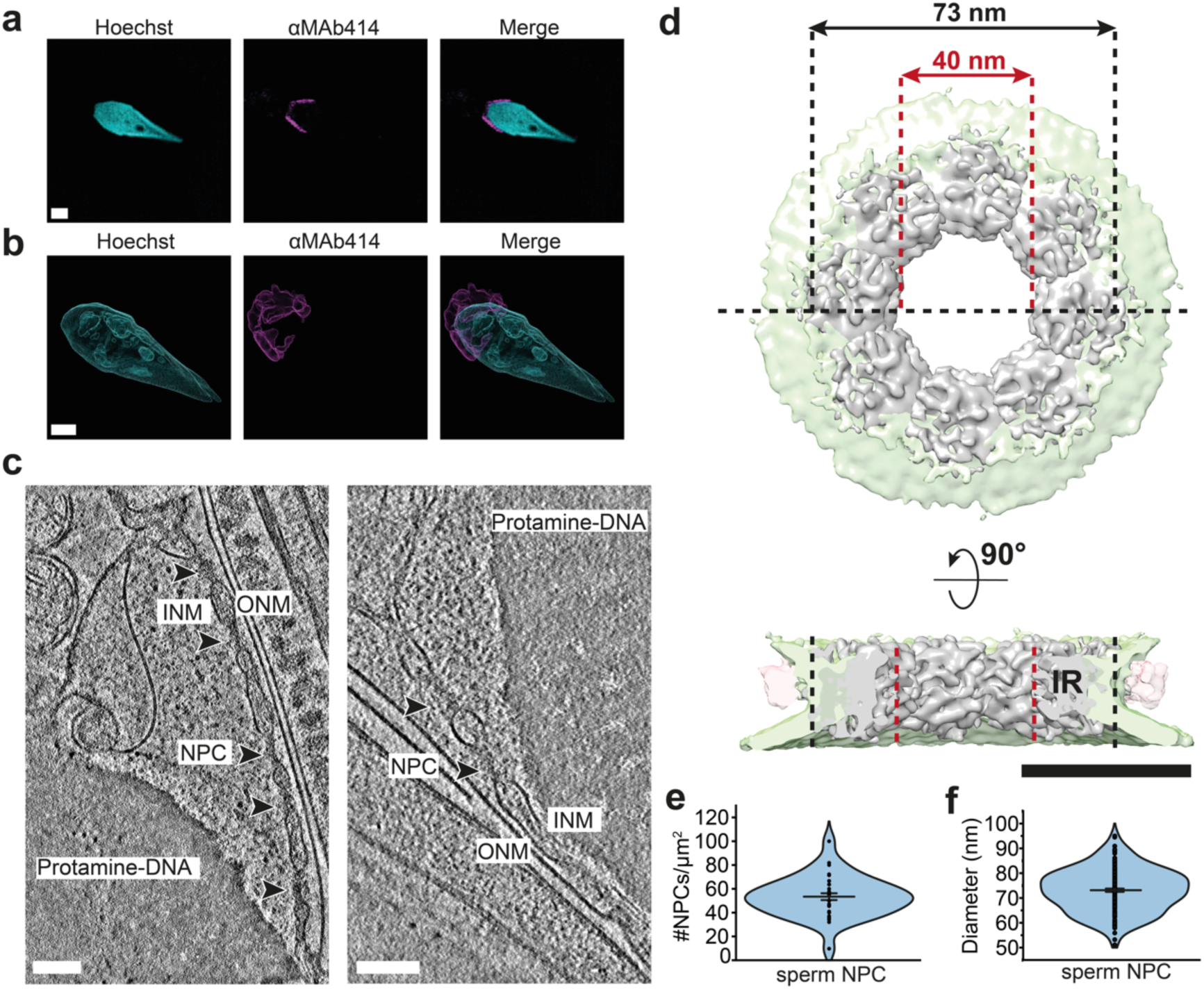
***In-cell architecture* of the human sperm NPC. a,** Structured Illumination Microscopy (SIM) of NPCs in human sperm cells. DNA stained with Hoechst shown in cyan and NPCs at the rear of the nucleus immunolabelled with antibody MAb414, shown in magenta. Scale bar = 1μm. Images are representative of data from more than ten biological replicates, with at least two technical replicates. **b,** 3D SIM reconstruction of the NPC region in the sperm cell shown in (**a**). Scale bar = 1μm. **c,** 2D slices of reconstructed tomograms of the NPC regions of two human sperm cells. Condensed DNA (Protamine-DNA), inner nuclear membrane (INM) and outer nuclear membrane (ONM) are shown. NPCs are indicated with arrowheads. Scale bar = 100 nm. Images show representative slices from hundred tomograms. **d,** *In-situ* structure of human sperm NPC – top view (above) and cross-sectional view (below). Inner Ring (IR) is shown in grey, nuclear membrane is shown in light green, luminal ring (LR) in light pink. Central channel diameter is indicated with red dashed lines, membrane-to-membrane diameter with dashed black lines. Scale bar = 50nm. **e,** Violin plot showing quantification of number of NPCs per nuclear membrane surface area in human sperm, from tomography data. Mean ± SE = 54 ± 3 NPCs/μm^2^, n= 34 sperm cells. **f,** Membrane-to-membrane diameter measurements of human sperm NPC from tomography data. Mean ± SE = 73.1 ± 0.7nm, n=174 NPCs from sixty tomograms.

A limitation of using fluorescence microscopy to study the organisation of protein complexes, is the lack of cellular context. To visualise individual human sperm NPCs in their native environment at higher resolution, we used cryo-ET^26^. Following vitrification of human sperm cells on cryo-EM grids, we used cryo-focused ion beam (cryo-FIB) milling for sample thinning. This was followed by tilt-series acquisition on the produced lamellae at NPC regions at the rear of the sperm head. Our reconstructed tomograms (Fig. 1c) showed the inner nuclear membrane (INM) and the outer nuclear membrane (ONM) with several NPCs stacked along a stretch of NE (Fig. 1c, arrowheads). Using this volumetric data, we measured NPC density in the targeted regions as 54± 3 NPCs/μm^2^ (Fig. 1e). This represents an approximately 7-fold increase in density relative to 5-10 NPCs/µm^2^ present at the NE^45^ of somatic cells, indicating a high degree of NPC clustering in human sperm cells.

### *In-situ* architecture of the human sperm NPC reveals the absence of cytoplasmic and nuclear rings and a highly constricted central channel

Next, we investigated the structure of the human sperm NPC by performing subtomogram averaging^10^, which resulted in a 3D density map of the native architecture of the complex (Fig. 1d and Extended Data Fig. 1d-g).

Unlike human somatic NPCs (Extended Data Fig. 1g)^2,20^, our *in-situ* average of the human sperm NPC showed that only the IR was present, and no signal for the CR, NR and nuclear basket was detected in the subtomogram average (Fig. 1d, Extended Data Fig. 1d, e and g, Supplementary Video 1). Furthermore, our cryo-ET data revealed a mean membrane-to-membrane NPC diameter of approximately 73nm and a central channel less than 40nm wide (Fig. 1d, f and Extended Data Fig. 1g). Compared to reported values of 92-105nm (membrane-to-membrane) and 54-60nm (central channel) for the *in-situ* NPCs in three different human cell lines^2,30,46^, and 82- 89nm (membrane-to-membrane) and 42-43nm (central channel) for the constricted *ex-cellulo* NPC^2,20^, our data indicates that the human sperm NPC adopts a highly constricted conformation at the NE *in-cellulo*.

Notably, our *in-situ* structure did not show densities accounting for the presence of the Y-complex (Fig. 1d, Extended Data Fig. 1d, e, g and Supplementary Video 1). To validate this, we performed immunoblotting and immunofluorescent labelling and SIM against the Y-complex component NUP107^10^ in sperm cells. Our data show that, in agreement with our map, NUP107 is absent in mature human sperm cells (Extended Data Fig. 2a and e).

We also tested for the presence of RanBP2 and TPR to validate the absence of the CR and NR, respectively, using immunoblotting and immunofluorescence labelling. In agreement with our structural data, both proteins were in very low abundance, and notably not co-localised with NPCs (Extended Data Fig. 2b, c). Moreover, the presence of both components could not be detected through immunoblotting of human sperm lysates (Extended Data Fig. 2f and g). Conversely, NUP93, a component of the IR, was present in large abundance and showed strong co-localisation with MAb414 (Extended Data Fig. 2d). As a control, all these proteins were detected in human fibroblasts and co-localised at the NE with MAb414 (Extended Data Fig. 3a-h).

Taken together, our structural, biochemical and imaging data reveal a reduced NPC scaffold architecture in mature human sperm cells, comprising exclusively the IR and displaying a constricted conformation.

### Reorganisation of NPCs during human spermatogenesis occurs following meiosis

To investigate at which stage of human spermatogenesis (Extended Data Fig. 3a) NPC reorganisation occurs, we performed immunohistochemistry (IHC) on approximately 5μm thick human testicular tissue obtained from five healthy donors. We used MAb414 as a marker for NPCs (Extended Data Fig. 3b and c, Extended Data Fig. 4a), peanut agglutinin (PNA) for acrosome labelling, and synaptonemal complex protein 3 (SYCP3) for identification of meiotic cells (Extended Data Fig. 3b and c). We distinguished cells at various differentiation stages according to these markers, their radial location within the seminiferous tubule, their shape and size. Our data show NPCs distributed uniformly around the NE in pre-meiotic spermatogonia (Extended Data Fig. 3c and Extended Data Fig. 4b), similar to somatic cells (Extended Data Fig. 1b, Extended Data Fig. 2). In cells undergoing meiosis (spermatocytes), MAb414 staining was also distributed around the whole nucleus, with some NPC-enriched regions, as previously reported using electron microscopy of thin plastic sections^47^. However, following meiosis, NPCs in round spermatids either localised at two opposing poles (Extended Data Fig. 4b) or at two flanking regions at the rear of the nucleus (Extended Data Fig. 4b; Extended Data Fig. 3c), suggesting that major reorganisation events take place at the final step of meiosis. Following the round spermatid stage, NPCs were found at their final location at the posterior side of the nucleus in both elongated spermatids and spermatozoa, opposite the acrosomal membrane (Extended Data Fig. 3c).

To further investigate NPC reorganisation during human spermatogenesis, we performed 3D rendering of our IHC confocal data from human testes sections (Extended Data Fig. 4b). This allowed us to measure the surface area/volume ratio of MAb414 signal at the NE, as a proxy of NPC clustering (Extended Data Fig. 4c). Our data show a drop in NPC surface area/volume ratio from 6.8μm^-1^ in spermatogonia to 5.8μm^-1^ in spermatocytes, and 3.60μm^-1^ in round spermatids, indicating sequential clustering of NPCs (Extended Data Fig. 4c). However, no significant change was detected between round spermatids and spermatozoa (3.89μm^-1^). These data suggest that clustering of NPCs is most pronounced during meiosis, and it has largely been achieved by the first stage of spermiogenesis (round spermatid).

To assess at which stage of spermatogenesis the CR and NR are no longer part of the NPC structure, we performed IHC against the Y-complex component NUP107 in human testes (Fig. 2a). NUP107 was present and co-localised with MAb414 at the NE of spermatogonia and spermatocytes (Fig. 2b). However, in post-meiotic cells NUP107 was either absent or no longer localised at the NE or at NPC regions identifiable with MAb414 (Fig. 2b arrowheads). A similar pattern was observed for RanBP2 (Fig. 2c and d) and TPR (Extended Data Fig. 3d and e) in tissue: both the proteins co-localised with MAb414 at the NE in spermatogonia and spermatocytes, but co-localisation was lost during spermiogenesis at round spermatid stage for RanBP2 (Fig. 2c and d -arrowheads) and elongated spermatid stage for TPR (Extended Data Fig. 3d and e - arrowheads). Conversely, NUP93 co-localised with MAb414 throughout spermatogenesis and in mature spermatozoa, as expected (Extended Data Fig. 5a and b).

**Fig. 2.**
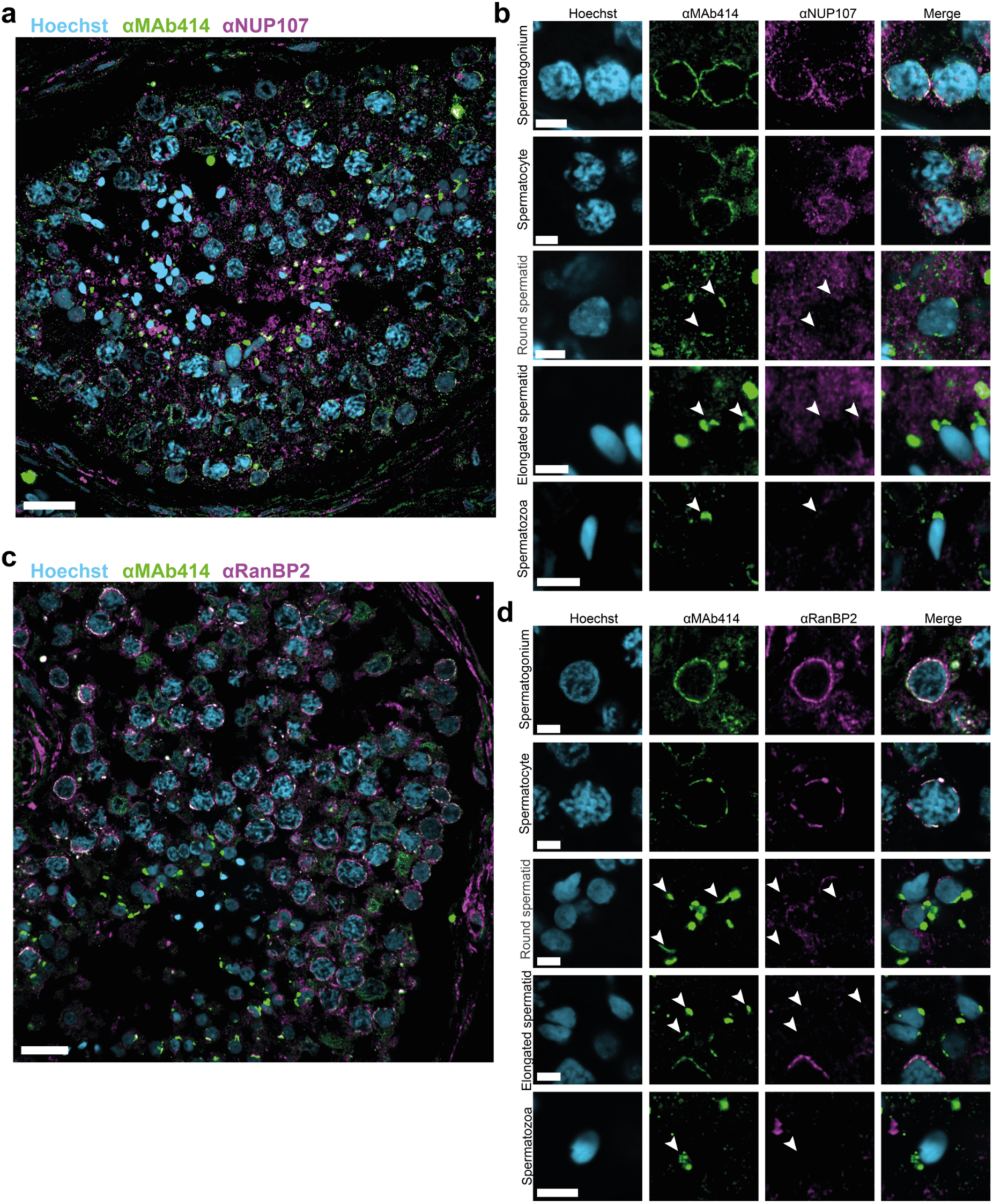
Absence of outer NPC components in post-meiotic human male germ cells. a, Immunohistochemistry and confocal imaging of a representative seminiferous tubule from healthy human testis tissue. DNA (Hoechst) is shown in cyan; NPCs (MAb414) in green and Y- complex component NUP107 is shown in magenta. Scale bar = 50μm. Image is representative of seminiferous tubules from five human donors, with at least two technical replicates. **b,** Detail of different cell populations within human seminiferous tubules, labelled as described in **(a)** showing re-distribution of NPCs at the NE in green. Arrowheads indicate loss of co-localisation between NUP107 (magenta) and MAb414 during differentiation. Scale bars = 5μm. **c,** Immunohistochemistry of human testis tissue showing the distribution of NPCs (MAb414, green) and cytoplasmic ring component RANBP2 (magenta) in a seminiferous tubule. DNA stained with Hoechst is shown in cyan. Scale bar = 50μm. Image is representative of seminiferous tubules from at least three human donors. **d,** Sub-populations of differentiating cells labelled as described in (**c**). Arrowheads indicate loss of co-localisation between RanBP2 and MAb414. Scale bars = 5μm.

Interestingly, during reorganisation of NPCs in the human tissue, the presence of strong NPC signal in the cytoplasm of different cell types was also detected. We used calreticulin as an ER marker in human testis sections and detected frequent co-localisation with cytoplasmic NPC signal at stages of differentiation between spermatogonia and elongated spermatids (Extended Data Fig. 5c and d, arrowheads). We postulate that cytoplasmic NPC signal in testes sections represents annulate lamellae^47^, which are often embedded in ER membranes^23^.

Altogether these data indicate that alterations to both the localisation and structure of NPCs during spermatogenesis occur after meiosis, when germ cells differentiate from spermatocytes to spermatids.

### Cryo-ET reveals the presence of filamentous structures embedded between the INM and ONM surrounding NPCs

Our tomographic analysis showed that NE regions adjacent to NPCs displayed intermembrane distances (distances between the INM and ONM) similar to those found in somatic cells, while these distances were much smaller in regions of the NE further from NPCs (NPC-distal region, Extended Data Fig. 6a). To quantify this, we performed morphometrics analysis^48^ of reconstructed tomograms (Extended Data Fig. 6a-c). Using these data, we calculated the modal value for each region^48^ and show that intermembrane distances significantly decrease from approximately 27nm in NPC-proximal regions to 9nm in NPC-distal regions (Extended Data Fig. 6b-d).

Interestingly, analysis of cryo-ET data collected at NPC-proximal regions revealed filamentous structures within the INM-ONM intermembrane space (Fig. 3a and b and Extended Data Fig. 6e). These structures were present in all tomograms collected at the NE of NPC-rich regions but absent in NPC-distal regions (Extended Data Fig. 6a, and d). Further characterisation and segmentation of the filamentous densities (Fig. 3a and Extended Data Fig. 6e) disclosed that they consisted of approximately 90 ± 0.2 nm in length (Fig. 3c). These filaments also displayed a high degree of bending and flexibility and were often found as two intertwined filaments of circa 12nm width (Fig. 3a, b, d, Extended Data Fig. 6e and Supplementary Video 2). Based on these measurements, we excluded actin filaments, microtubules, lamins, vimentin and other cytoplasmic intermediate filaments^49^ as possible sources for our filamentous structures.

**Fig. 3.**
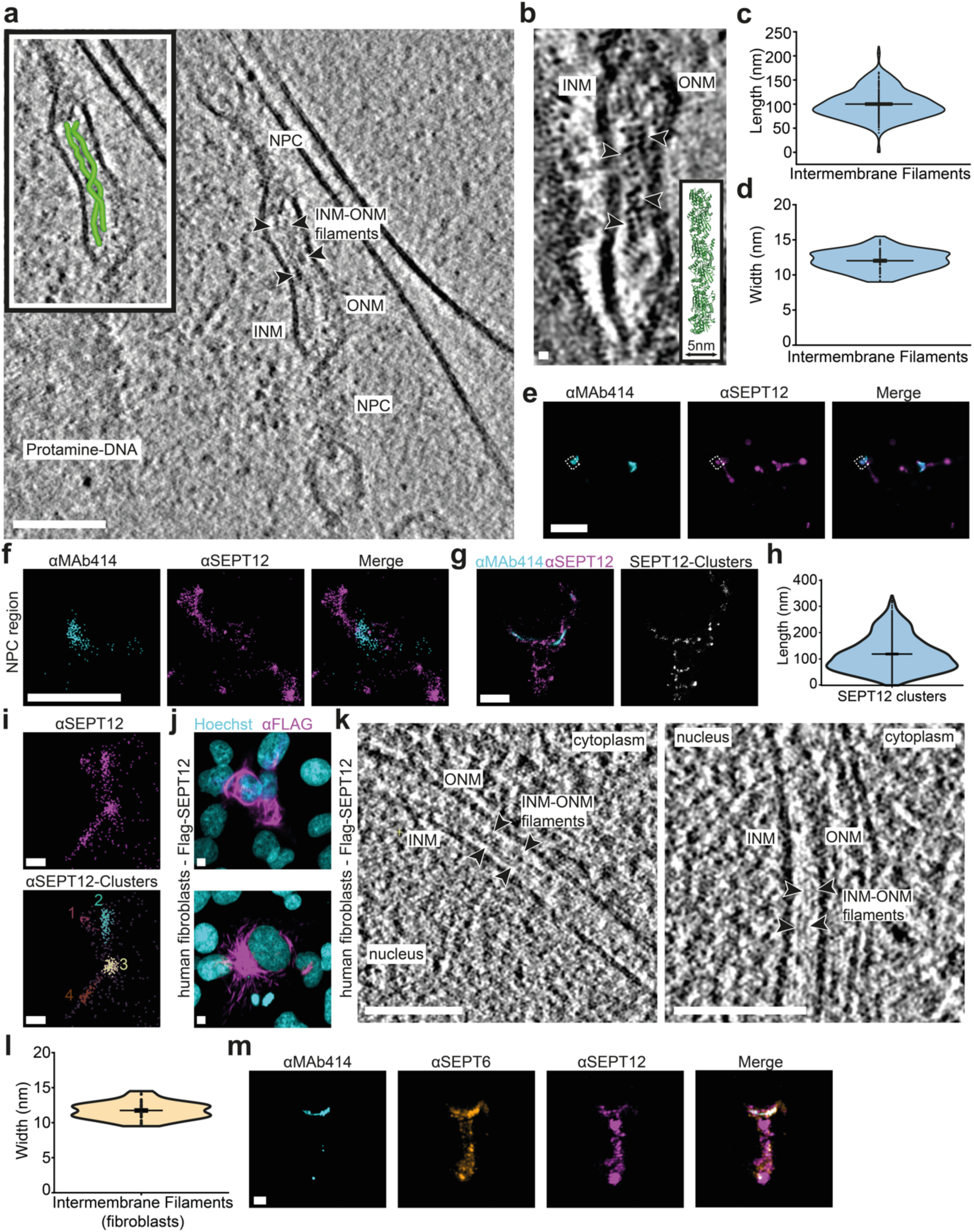
Filamentous structures within the INM and the ONM space in human sperm cells. a, Cryo tomographic slice showing the INM and ONM of human sperm cells. Scale bar = 100nm. Arrowheads indicate filamentous structures observed in the INM-ONM space. Segmentation of intermembrane filaments shown in green (top-left corner inset). Image is representative of cryo tomographic collected across three biological replicates (donors). **b,** Additional example of nuclear envelope region with intermembrane filamentous structures indicated with arrows showing width of approximately 6.5nm; scale bar = 10nm. Inset detail shows structure of a SEPT2-SEPT6-SEPT7 hetero-oligomer (PDB:9bht). Inset scale bar = 5nm. **c,** Length measurements of INM-ONM intermembrane filaments calculated from cryo tomographic data. Mean ± SE = 90.9 ± 0.2nm, n = 187 filaments from three biological replicates (donors) and fifty tomograms. **d,** Width measurements of INM-ONM filaments (dimeric form) from cryo tomographic data. Mean ± SE = 12.0 ± 0.2nm, n = 63, from three biological replicates (donors). **e,** Wide-Field imaging of human sperm cells immunofluorescently labelled with SEPT12 (magenta) and MAb414 (cyan). White rectangle shows NPC region selected for super-resolution imaging STORM, shown in (**f**). Scale bar = 10μm. **f,** two-colour STORM of NPC region of the human sperm cell shown in (**e**). Scale bar = 1μm. **g,** two-colour STORM of whole human sperm cell showing SEPT12 (magenta) localisation at the annulus, neck and NPC region co-labelled with MAb414 (cyan). Right panel shows SEPT12 clusters (white) generated with HDBSCAN analysis. Scale bar = 1μm. **h,** Violin plot showing length measurements of SEPT12 clusters in human sperm cells from STORM data and HDBSCAN analysis. Mean ± SE = 118nm ± 2.5nm, n = 739 clusters. **i,** Zoom- in showing four examples of SEPT12 clusters (numbering 1-4). Left panel shows original STORM data and right panel shows superimposition of clusters detected with HDBSCAN. Scale bar = 100 nm. **j,** confocal imaging of human fibroblasts transfected with Flag-SEPT12 immunolabelled with anti-FLAG antibody (magenta). DNA is shown in cyan (Hoechst). Scale bar = 5μm. **k,** cryo tomographic slices of human fibroblasts transfected with FLAG-SEPT12. Images show the presence of filamentous structures within the intermembrane space of the NE (arrowheads). Scale bar = 100nm. **l,** width measurements of INM-ONM filaments (dimeric form) from cryo tomographic data of human fibroblasts transfected with FLAG-SEPT12. Mean ± SE = 11.7nm ± 0.2nm, n = 29 filaments. **m,** SIM image showing immunofluorescently labelled human sperm NPCs (MAb414, in cyan), SETP12 (magenta) and SEPT6 (orange). Scale bar = 1μm. Results were validated in more than three independent experiments.

Previous work on mammalian sperm highlights the importance of septins - polymerising GTP-binding proteins - in spermatogenesis and male fertility^50^. For example, septin 12 (SEPT12) is expressed exclusively in testes and specific mutations or changes in its expression levels lead to infertility due to defects in the sperm annulus, or nuclear damage and maturation arrest at the spermatid stage^51–53^. In sperm, SEPT12 has also been shown to interact with NE proteins, including Lamin B1 (LB1), B2/B3 (LB2/3)^54^, and NDC1^55^, a membrane component of the NPC. Additionally, structural characterisation of septins shows their organisation as approximately 5nm-wide single filaments^56^ (Fig. 3b inset), and 11nm intertwined filaments^57^ corresponding to the dimensions of the filamentous densities at NPC-rich regions in our tomograms (Fig. 3a, b and d and Extended Data Fig. 6e). Therefore, we considered SEPT12 as a plausible candidate to investigate the nature of the intermembrane filaments.

Firstly, we performed immunoblotting against SEPT12 confirming, as expected, its presence in human sperm lysate (Extended Data Fig. 6f). Previous studies have shown localisation of SEPT12 at the neck and annulus of human sperm cells^58,59^, however, our wide-field immunofluorescence data showed SEPT12 signal also extending its localisation to the NPC-proximal region of the NE (Fig. 3e). To obtain higher resolution detail of SEPT12 organisation in this region, we performed Stochastic Optical Reconstruction Microscopy (STORM). STORM imaging revealed SEPT12 co-localisation with NPCs immunolabelled with MAb414 (Fig. 3f and g), consistent with our cryo-ET data (Fig. 3a and Extended Data Fig. 6e). Next, we measured the length of SEPT12 filaments imaged with STORM and determined that SEPT12 forms elongated clusters with a mean length of 118 ± 2.5 nm (Fig. 3h and i). These measurements were similar to the ones obtained from cryo-ET data (Fig. 3c) and the discrepancy might be explained by the lower resolution of STORM data (approx. 30nm) and the additional presence of antibodies.

To provide further evidence in support of SEPT12 as the filamentous protein localised within the inter-membrane space of the NE, we transiently expressed FLAG-SEPT12 in human fibroblasts (Fig. 3j). We performed cryo-FIB-milling on these cells and acquired tilt-series on the produced lamellae. Our tomographic analysis revealed presence of ∼12nm intertwined filaments (Fig. 3k and l) within the nuclear intermembrane space, absent in control fibroblasts (Extended Data Fig. 6h). These data strongly support our hypothesis that Septins are the source of the INM-ONM filaments in sperm cells.

Septins have been reported to form both homo and hetero oligomers with other septin isoforms^56,60^. In sperm, SEPT12-SEPT7-SEPT6-SEPT2 and SEPT12-SEPT7-SEPT6-SEPT4 hetero-oligomers have been hypothesised based on co-immunoprecipitation studies and due to their co-localisation at the neck and annulus regions^61^. To investigate the possibility of additional septin isoforms being present at the NE of NPC-rich regions, we used immunoblotting against SEPT6 in human sperm lysate (Extended Data Fig. 6g) and performed SIM on immunofluorescently labelled human sperm cells. Our imaging data showed that SEPT6 and SEPT12 co-localised in the neck and annulus regions of sperm cells, as expected, but also at the NPC region (Fig. 3m). Higher-resolution imaging with STORM also showed partial co-localisation of SEPT6 and SEPT12 in human sperm cells, including at the NPC region (Extended Data Fig. 6i and j), suggesting that mixed populations of SEPT12 hetero- and homo-oligomers might be present at the NPC-rich NE regions. This also confirms the previously described heterogeneity of septin filaments^56,60,61^.

As previously mentioned, interactions between SEPT12 and lamin isoforms have been reported in sperm cells. We also found the presence of lamin isoforms in NPC-proximal regions of our reconstructed tomograms (Extended Data Fig. 7a). Immunoblotting of sperm lysates against LB1, LB2 and Lamin A/C (LMNA/C) confirmed their presence in human sperm (Extended Data Fig. 7b). To investigate the cellular localisation of the different lamin isoforms, we performed immunofluorescence labelling, coupled to SIM imaging in human sperm cells. Both LB1 and LB2/LB3 were located at the rear of the nucleus, co-localising with MAb414 (Extended Data Fig. 7d and e). Conversely, LMNA/C appeared distributed around the whole NE (Extended Data Fig. 7c).

Overall, according to our findings, we propose that the filaments found within the INM-ONM space are homo- and hetero-oligomers of septin isoforms.

### Nuclear transport is perturbed in human sperm cells

Our *in-situ* reconstruction of the human sperm NPC revealed large architectural changes, including the absence of the CR and NRs, and a highly constricted central channel. To understand how these changes affect NPC function, we investigated potential disruptions to Ran-dependent active transport and nuclear diffusion into the nucleus.

The maintenance of a Ran-gradient in the cell depends on several factors, including RanGAP1, the Ran GTPase activating protein, which is mostly localised on the cytoplasm and at the outer surface of the NE of mammalian somatic cells^62^ (Extended Data Fig. 8a). We investigated the presence and localisation of both Ran and RanGAP1 in human sperm using immunofluorescence and immunoblotting. We detected low levels of Ran almost exclusively localised to the neck and cytoplasmic region of sperm cells^40^ (Fig. 4a and d). This differed to its localisation in human fibroblasts, where it was highly enriched in the nucleus during interphase (Extended Data Fig. 8a and d). Conversely, we could not detect RanGAP1 through immunoblotting in sperm lysates, and immunofluorescence data showed very low signal (Fig. 4b and e) compared to fibroblasts (Extended Data Fig. 8b and e). We then tested the presence and localisation of the main nuclear transport receptor importin **β**1. In human fibroblasts, as expected, importin **β**1 was present and mostly nuclear (Extended Data Fig. 8c and f), whilst in human sperm cells, we could only detect reduced levels of importin **α**1^40^ (Fig. 4c and f), localised exclusively to the outside the nucleus in the neck region (Fig. 4c). Taken together, these data point towards a disrupted active transport in human sperm cells.

**Fig. 4.**
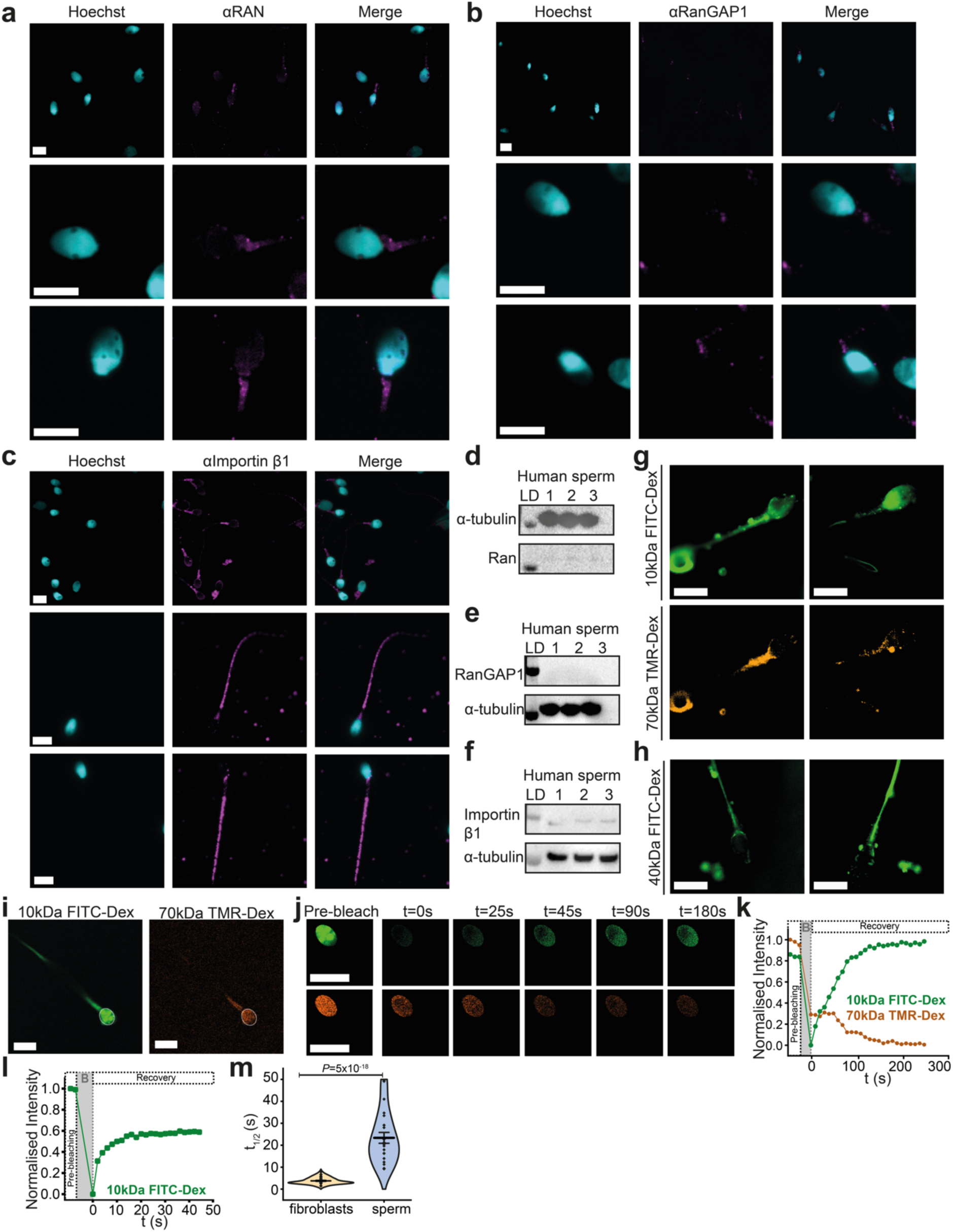
Nuclear active transport in human sperm is perturbed and passive diffusion is reduced. a, Immunofluorescence labelling of human sperm cells showing Ran. Scale bar = 5μm. Results were validated in more than three independent experiments (technical replicates). **b,** Immunofluorescence labelling of human sperm cells showing RanGAP1. Scale bar = 5μm. Results were validated in more than three independent experiments. **c,** Sperm cells immunolabelled with Importin β1 antibody. Scale bar = 5μm. Results were validated in more than three independent experiments. **d-f,** Western blots against **d,** Ran, **e,** RanGAP1 **f,** and Importin β1 using human sperm lysate. α-tubulin was used as loading control and three replicates are shown. **g-h,** Diffusion test in human sperm cells using digitonin for the delivery of fluorescently labelled dextrans of different molecular weights. Scale bars = 5μm. **g,** two human sperm cells with both 10kDa-FITC Dextran and 70kDa-TMR Dextran showing exclusive passive diffusion of 10kDa dextran to the nucleus after digitonin treatment. Data are representative of at least three independent experiments. **h,** Human sperm cell showing inability of 40kDa-FITC dextran to passively diffuse into the nucleus, following digitonin treatment. Results were validated in more than three independent experiments. **i,** Sperm cell showing even distribution of 10kDa-FITC and 70kDa-TMR dextrans following electroporation. Scale bars = 5μm. Bleaching region used for Fluorescence Recovery After Photobleaching (FRAP) shown as a white-dotted circle. **(j)** Fluorescent images showing the recovery time-course for the cell shown in (**i**). Scale bars = 5μm. (k) FRAP recovery curve for both 10kDa-FITC and 70kDa-TMR Dextran in cell showing in (I and J). (l) Example of a recovery curve for a human fibroblast electroporated with 10kDa FITC dextran. (m) Calculated half times of recovery (t1/2) for 10kDa-FITC FRAP experiments in human fibroblasts and human sperm cells. Mean ± SE: t1/2(fibroblasts) = 3.8 ± 0.2s, n = 50 cells; t1/2(sperm)=23.4 ± 2.6s, n = 23 cells. Statistical Analysis: t- test, *P*-value is shown.

Previous *in-situ* structural work in *S. pombe* showed that NPC constriction, following hyperosmotic shock, can affect passive diffusion rates of small proteins into the nucleus^31^. As our cryo-ET data showed a highly constricted NPC (Fig. 1d, f and Extended Data Fig. 1d and g), with a central channel volume approximately 50% smaller than in somatic cells^30^, we also tested this hypothesis. For this, we treated sperm cells with digitonin, as it can permeabilise the cytoplasmic, but not the nuclear membrane of mammalian cells, to deliver fluorescently labelled dextrans of different molecular weights (10kDa, 40kDa and 70kDa). We detected 10kDa-FITC dextrans in the sperm nucleus due to passive diffusion, but we could not detect the presence of 40kDa-FITC or 70kDa-TMR dextrans in the nucleus (Fig. 4g and h). The same results were obtained in human fibroblasts (Extended Data Fig. 8g-i).

After ascertaining that 10kDa dextrans could freely diffuse into the nucleus of sperm cells, we compared passive diffusion rates between human sperm cells and fibroblasts using Fluorescence Recovery After Photobleaching (FRAP). First, we electroporated human sperm cells with both 10kDa-FITC and 70kDa-TMR dextrans to deliver both fluorescent molecules to the whole cell, including the nucleus (Fig. 4i). Then, following bleaching of both fluorophores in the nuclear region, we detected fluorescence recovery of 10kDa-FITC but not of 70kDa-TMR dextran, as expected and also indicative that the nuclear membrane remains intact after electroporation (Fig. 4j and k). We then used 10kDa-FITC dextrans for all passive diffusion measurements in sperm cells and fibroblasts (Fig. 4k-m). Our FRAP measurements in human sperm cells displayed higher variability and a large increase in half-time of recovery, relative to human fibroblasts (mean ± SE: sperm t1/2= 23.4 ± 2.5 s; fibroblasts t1/2= 3.8 ± 0.2s; Fig. 4m). Although other variables might also contribute to this large reduction in passive diffusion rates in sperm cells, we hypothesise that NPC constriction is the main causes of this^31^.

Overall, our data suggest a direct link between the reduced, constricted NPC architecture and perturbation of both Ran-dependent active transport and passive diffusion to the nucleus in human sperm cells (Fig. 5).

**Fig. 5.**
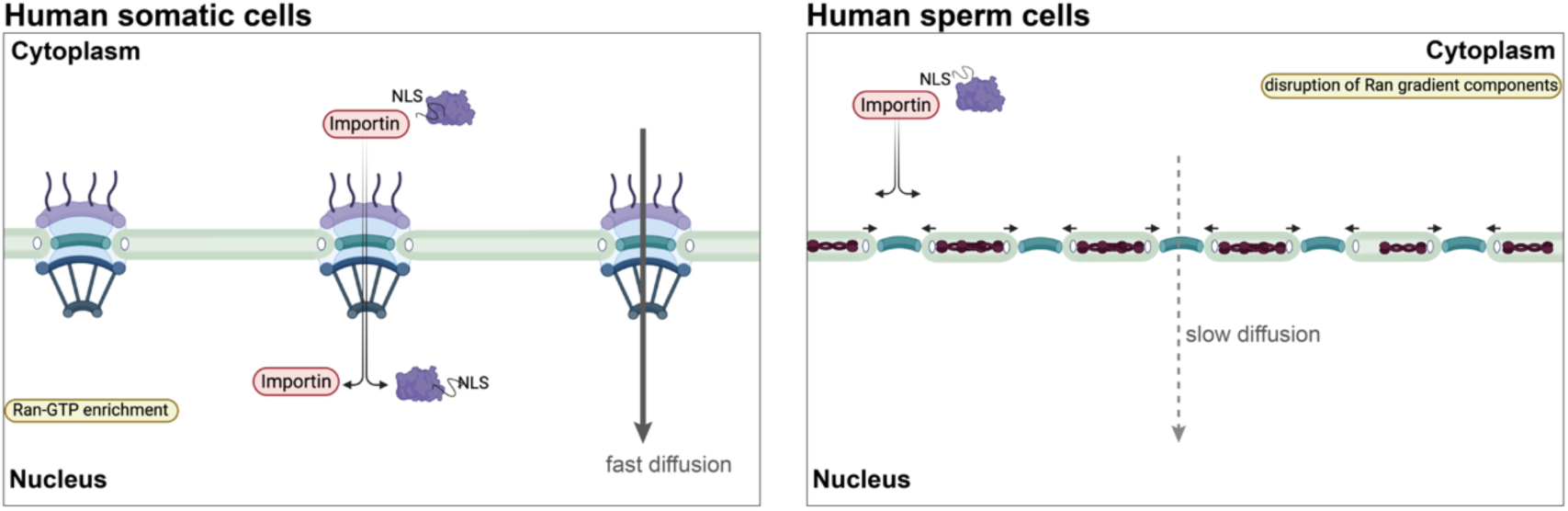
Structural and functional changes to the NPC of human sperm cells. Human sperm NPCs have a reduced scaffolding, showing absence of the Y-complexes and outer rings, and a highly constricted central channel (indicated by arrows, right panel). These structural changes result in slow passive diffusion of small molecules into the nucleus and in disruption of the Ran- dependent active transport.

### Discussion

In the last decades, specific roles of NPCs in tissue development and differentiation have been revealed^63^. Despite this, structural data are lacking behind. In this study, we investigated the architecture and the function of the human NPC in male germ cells using an integrative *in-situ* structural biology approach.

We describe major changes to the NPC structure in human sperm cells, relative to somatic cells^2^, including loss of the outer rings (CR and NR), and a high degree of constriction of the central channel (Fig. 1, Extended Data Fig. 1 and 2).

In somatic cells, Y-complexes are one of the main scaffold components needed for the assembly of NPCs^10^. However, densities for the Y-complex were not present in our *in situ* EM structure (Fig. 1d, Extended Data Fig. 1d, e and g and Supplementary Video 1). Consistent with this, we could not detect the Y-complex subunit NUP107 in human sperm lysates through immunoblotting, and immunofluorescence revealed only low signal not localised at NPCs (Extended Data Fig. 2a and e). Our data are further validated by previous proteomics studies^64^ that did not detect NUP107 in mature human sperm. Despite the absence of the Y-complex, NPCs are highly clustered at the caudal region of the NE (Fig. 1a and b, Extended Data Fig. 1c and Extended Data Fig. 4b and c), with an approximately seven-fold increase in NPC density, relative to previously reported values for human somatic cells^45,65^ (Fig. 1e).

Our human tissue data show that NPC clustering was most pronounced as cells exited meiosis and entered spermiogenesis. In round spermatids, NPCs no longer uniformly decorated the nucleus as they did in spermatogonia, but instead localised exclusively at two opposing lateral poles (Extended Data Fig. 4b and Extended Data Fig. 3c). This is consistent with previous evidence from electron microscopy images of thin plastic sections in rodents and humans, showing accumulation of NPCs at the caudal end of the nucleus of late-stage spermatids^41,66^. It is possible that clustering of NPCs at the NE reaches its maximum at the exit of meiosis and for the remaining steps of spermiogenesis nuclear membranes enriched with NPCs descend to the caudal region of the head. Further studies are needed to understand how relocation of NPCs occurs, but biochemical evidence in rats suggests a microtubule-dependent movement mediated by interactions between the IR and the kinesin molecular motor KIFC1 in post-meiotic spermatids^5^.

Previous RNAi work in *D. melanogaster* spermatogenesis demonstrated the crucial role of the IR in initiating meiosis^4^, whilst suggesting that the CR, NR and Y-complexes are dispensable for this process^67^. Our human tissue imaging data agree with these findings, in fact we show that the loss of the outer NPC rings starts during meiosis, concurrently with NPC clustering and DNA hyper-condensation. In human testes, the Y-complex component NUP107 colocalised with MAb414 in both spermatogonia and spermatocytes but was lost in round spermatids (Fig. 2a and b). RanBP2, usually at the CR, was also no longer found to co-localise with MAb414 in round spermatids and the signal was mostly lost in elongated spermatids and spermatozoa (Fig. 2c and d). Similarly, TPR, a nuclear basket component, lost co-localisation with NPCs at the spermatid stage (Extended Data Fig. 3d and e). Previous biochemical work showing the absence of NUP153, another nuclear basket component, in the testis of adult rats^5^, further validates our findings. Interestingly, NUP107, RanBP2 and NUP153 are present in oocytes^23^, suggesting possible complementary roles of NPC components post-fertilisation. Pre-assembled IRs delivered by sperm cells during fertilisation may facilitate reincorporation of the remaining nucleoporins^14,15,34^, and these may aid male pronuclear movement^44^.

Another major finding of this study, enabled by *in situ* cryo-ET, was the unexpected presence of filamentous structures embedded in the intermembrane space between the INM and ONM (Fig. 3a-l, Extended Data Fig. 6e and Supplementary Video 2). We propose that these filaments are oligomeric forms of septins, in particular of SEPT12 a testis and lymphocyte-specific isoform^68^ – and SEPT6 (Fig. 3e-l and Extended Data Fig. 6e-j). SEPT12 is essential for successful spermatogenesis, and its depletion or mutation is known to cause defects in sperm cell morphology and lead to infertility^51–53,61^. However, it is not clear what its specific role during differentiation is. In somatic cells, the role of septins is also not well understood, but evidence suggests they form apolar filaments that interact with phospholipids and restrict diffusion of membrane-bound proteins^57,69^. Septins are also thought to have a role in maintaining cell cortex rigidity at specific sites and in subcellular compartmentalisation^70,71^. Previous studies in sperm cells show that SEPT12 interacts with different NE components including lamins (LB2/LB3, LB1), the LINC component SUN4, and NDC1^54,55,72^, a membrane component of the NPC. Although previous reports only detected SEPT12 at the annulus and neck of sperm cells^54,55,61^, our super-resolution imaging data showed its presence extending to NPC regions (Fig. 3f and g, Extended Data Fig. 6i and j), in agreement with the location of these filaments in cryo-ET data. Following, over- expression of SEPT12 in human fibroblasts, our cryo-ET data clearly showed the presence of intertwined filaments approximately 12nm wide, within the intermembrane space of the NE (Fig. 3j-l), validating our hypothesis. Our conclusions are also in line with the localisation of known NE interactors of SEPT12, including NDC1, and LB1, LB2/LB3, which we show to be present at the INM side of NPC-rich regions (Extended Data Fig. 7).

SEPT12 oligomerisation at the NE will, most likely, affect the local mechanics of the membrane^57,70^. This is important, as NPC dynamics respond to local forces altering membrane tension, such as cytoskeletal filaments acting on the NE^31,32,35,36,73^. Our *in-situ* model for the sperm NPC shows a highly constricted central channel. Indeed, the central channel diameter was measured at ≤40nm, compared to approximately 55- 60nm in human somatic cells (Fig. 1d and Extended Data Fig. 1g). Lack of scaffold NUPs should lead to loss of structural rigidity of NPCs and promote the expansion of the central channel, as previously reported^30,31,35^. Our reduced NPC structure is, instead, constricted. We propose that this is the result of forces introduced by the presence of the septin network found within the nuclear inter-membrane space. Septin filaments are in fact known to be important in stabilising membranes, by introducing additional rigidity^70^. As a result, they may prevent NE instability or rupturing events during sperm cell movement and fertilisation. Further studies on septins role in sperm cells are still required to answer these questions.

Here, we also propose that functional changes accompany the structural differences found in the sperm NPC. For example, RanBP2, crucial for mRNA export^74^ and nuclear import^21^, largely lost co-localisation with MAb414 in human tissue following meiosis and could not be detected in mature sperm (Fig. 2c and d). Similarly, the absence of basket NUPs could affect nucleocytoplasmic transport by changing the availability of docking sites for transport receptors^75^. In line with this, we showed that in mature sperm cells, the Ran activator protein, RanGAP1, also interactor of RanBP2^21,62^, could not be detected using immunoblotting (Fig. 4e) and immunofluorescence showed very low signal in the tail region (Fig. 4b). Additionally, whilst we detected the presence of low levels of Ran and Importin **α**1, we also showed their enrichment in the sperm tail (Fig. 4a, c, d and f), contrasting somatic cells, where their signal is largely nuclear^76^ (Extended Data Fig. 8a, c, d and f). Ran, in particular, was localised to the centrosomal neck region (Fig. 4a), which supports its proposed function in microtubule assembly, as reported in rat spermatids^77^. In addition, our structural data also showed absence of the nuclear basket, and our light microscopy analysis showed reduced levels of TPR, not colocalised with MAb414 in human sperm cells (Fig. 1d; Extended Data Fig. 2c). An exclusively cytoplasmic Importin **α**1 would, in fact, prevent the import of TPR or NUP153 into the nucleus^78^ and, simultaneously, the unavailability of nuclear basket components would disrupt Importin **α**1-mediated transport^18^. Hence, overall, we accumulated evidence that suggests a disruption of Ran-dependent active transport in human sperm cells, which a recent study proposes playing a critical role for the assembly of the outer NPC rings^24^.

Due to the large reduction in the NPC central channel diameter in sperm cells (Fig. 1d and f; Extended Data Fig. 1g), we also hypothesised that there would be changes in passive diffusion rates to the nucleus. *In situ* structural work of the *S. pombe* NPC^31^ has previously shown a strong correlation between NPC diameter reduction and diffusion rates of small molecules into the nucleus. In agreement with this, our data showed an approximately six-fold increase in half-times of recovery for small fluorescently tagged dextrans in the nucleus of human sperm cells, compared to human fibroblasts (Fig. 4i-m).

Overall, our study reveals the architecture of the NPC in human sperm cells solved using a state-of-the-art *in-situ* integrative structural biology approach. A reduced and constricted NPC architecture represents a plausible mechanism to limit nuclear transport in differentiated sperm cells, which have to safeguard the paternal DNA while delivering the whole nucleus to the oocyte^44^. Furthermore, our findings showing the exclusive presence of the IR in mature sperm cells support a conserved function for the IR in meiotic progression, as shown in *D. melanogaster*^4^, and in NPC relocation, as shown during rat spermiogenesis^5^.

Our work emphasises how NPC compositional variability allows functional adaptation^19,79,80^ and underlines the crucial regulatory importance of NPCs in spermatogenesis and animal development^63^. We expect electron-cryo-tomography to be pivotal in discovering changes in NPC architecture in the context of different human cells and tissues. In perspective, specialised cells, such as sperm cells, may have additional compositional rearrangements of macromolecules, necessary for their function, and a variety of quaternary structures may be a common feature of gametes.

## Methods

### Human sperm and testis tissue samples

Human mature sperm cells were purchased from the European Sperm Bank (https://www.europeanspermbank.com/en) which sent us sperm cells from 25 random healthy men. The samples had been previously processed, tested by the bank according to the requirement of the national competent authorities, GMP and WHO standards, as well as EU directives: 2006/86/EC, 2006/17/EC, 2004/23/EC, 2015/565. Sperm cells were screened for morphology, genetic diseases, blood-borne viruses and motility. Consent to use the samples for research purposes was given to the European Sperm Bank.

Human testis FFPE sections (5-10μm thick) from five different individuals, aged 18-55 years of age and with no history of infertility, were obtained from the Imperial College Healthcare Tissue Bank (Department of Surgery and Cancer). All the experimental procedures were approved by the Imperial College Healthcare Tissue bank (Research Ethics Committee approval numbers: 22/WA/0214; ICHTB HTA license: 12275).

Human sperm straws were thawed to room temperature and used immediately after thawing. Cells were checked for motility under an optical microscope before grid preparation or other experiments.

### Vitrification

EM grids (Quantifoil R 1/4 Au 200 mesh, Quantifoil Micro Tools GmbH) were glow discharged with an Edwards S150B glow discharger at 30mA for 45sec on both sides of the grid. Plunge freezing was carried out with a Leica GP2 or with a custom-made manual plunger using a sperm concentration of around 3x10^7^ cells/mL and 3.5µl per grid. In some cases, multiple applications of 3.5µl were used to increase the density of cells per grid square. Blotting time was 6-9 sec and liquid ethane was used as the cryogenic.

### Cryo focused ion beam (FIB) milling

CryoFIB was carried out using three different instruments. Manual milling was performed using a Scios cryoFIB/SEM (Thermo Fisher Scientific), and automatic milling using either an Aquilos II cryoFIB/SEM (Thermo Fischer Scientific) or a Crossbeam 550 (Zeiss GmbH). Milling was performed according to previously published protocols^81,82^. Briefly an inorganic platinum layer (∼50nm) was sputter- coated in a Quorum chamber (Scios and Crossbeam 550) or in the main chamber (Aquilos II). Following this, a gas injection system (GIS) was used to cover the cells with a layer of organometallic platinum (Trimethyl [(1,2,3,4,5-ETA.)-1 Methyl-2, 4- Cyclopentadien-1-YL] Platinum) to protect cells from curtaining during milling. A milling angle of 7-8°, relative to the grid’s plane, was selected for milling. Milling was carried out stepwise using a staircase pattern (0.5nA, 0.3nA, 0.1nA, 30pA) up to 10µm lamella width and 150nm final lamella thickness.

### Tilt series acquisition and tomogram reconstruction

Tilt series were collected on a 300kV Titan Krios G3i transmission electron microscope (Thermo Fisher Scientific) equipped with a Bioquantum energy filter (Gatan) and a K3 direct electron detector (Gatan). Images were acquired with a pixel size of 1.63Å using Serial EM PACE tomo scripts^83^ and a dose-symmetric tilt scheme^84^ with 3° increments. The total electron dose was 100e/Å^2^ with each tilt 3 e/Å^2^ split in a movie of 6-12 frames, and defocus -2.5 to -5µm.

All movies were imported into Warp^85^ for gain and motion correction, tilt selection, CTF correction. Tomograms were reconstructed after aligning the tilt series with AreTomo^86^; Etomo^87^ was also used in difficult cases.

### Subtomogram averaging

NPC averaging was performed as described previously^29^ with modifications. Specifically, 338 NPC volumes were picked using Imod^87^ using two points for each NPC to give them a direction towards the cytoplasm. Model files were converted into a star file for Warp 1.1.0 extraction using a custom script^88^. After extraction at bin 6 (9.78Å/pix) volumes were imported into RELION 3.1^89^. An initial model was obtained with relion_reconstruct and volumes were classified (3 classes, T=1, no alignment) and refined using the RELION pipeline. The resulting NPC map was then split into the eight spokes using napari-subboxer^90^ (https://github.com/alisterburt/napari-subboxer) and the resulting ∼2600 subvolumes were used for 3D classification to discard bad particles (5 classes, T=1) and consensus refinement. Ellipsoidal masks were created in Dynamo^91^ to carry out focused refinement for the cytoplasmic region, the inner ring region, the nucleoplasmic region and the luminal region of the NPC spoke^29^. Using dynamo2m^92^ subvolumes were exported to M^93^ and refined for 3 sub-iterations using one at a time ‘Image warp grid, particle poses, stage angles and volume warp grid’. Final refinements were performed in RELION (Extended Data Fig. 1f).

UCSF Chimera was used to visualise maps, to create a composite map of the whole C8 symmetric NPC using Chimera’s “*fit-to-map*” for each of the single refined spokes, and to make figures and the Supplementary Video 1.

### Tomogram segmentation and analysis

The nuclear membrane was segmented manually in Imod^87^ using Drawing Tools and linear/meshing interpolation. Filaments in the nuclear membrane were traced using Imod (https://www.andrewnoske.com/wiki/IMOD_-_segmenting_tubes) which also computes their length and their width was measured manually. NPC diameters were measured manually in Imod and their density per µm^2^ was calculated dividing the total nuclear membrane surface area from 35 tomograms by the NPCs number. NPC central channel volume was estimated using only the IR central channel volume calculated as a cylinder with a height of 20nm as in^30^. The distance from the inner to the outer nuclear membrane was estimated using surface morphometrics^48^.

### Cell culture

Human skin fibroblasts derived from AG10803 and immortalised with SV40LT and TERT were a gift from Delphine Larrieu. Cells were cultured in Dulbecco’s modified Eagle’s medium, supplemented with 10% foetal bovine serum (FBS) and penicillin/streptomycin. Cells were maintained at 37°C, 5% CO2.

For digitonin experiments, sperm cells or fibroblasts were incubated for 30 minutes with 2μg/mL digitonin and 5μM dextrans (10kDa-FITC, 40kDa-FITC or 70kDa-TMR dextrans) at 37°C, 5%CO2. Cells were then washed with PBS and allowed to recover for 1h before fixation and imaging. For Hoechst staining, live cells were incubated for 10min with Hoechst 33342 (Thermo Fischer Scientific) at 37°C, 5%CO2 and then washed with PBS before fixation or imaging.

For mammalian cell transfections, human fibroblasts were seeded on glass coverslips #1.5 or tissue culture 6-well dishes and left to adhere overnight. FLAG-SEPT12 (pcDNA3.1 FLAG-SEPT12 plasmid, Invitrogen) was then transiently transfected using Lipofectamine 3000 (Invitrogen) for 48h. For immunofluorescence experiments, cells were fixed and labelled as described in ‘Immunofluorescence’. For electron-cryo- tomography experiments, cells transfected in 6-well plates were detached after 48h using 0.25% Trypsin-EDTA (Gibco) and seeded on EM grids pre-functionalised with 50μg/mL of fibronectin bovine plasma (Sigma). Cells were then left to attach for 5 hours at 37°C, 5% CO2 before sample vitrification and cryo-FIB milling.

### Antibodies

Primary antibodies for NPC components were selected based on their antigenicity to ensure minimal cross-reactivity with other NUPs. For antibody details and dilutions used see Extended Data Table 1.

### Immunofluorescence

Human fibroblasts were seeded on #1.5 glass coverslips for 24h at 37°C, 5% CO2 before fixation and immunofluorescence procedures. Human sperm cells were spread on #1.5 coverslips pre-functionalised with Poly-L-lysine (PLL) and left at room temperature until dry. Cells were then fixed for 10 minutes at room-temperature (RT) in 4% PFA, then washed in PBS. Samples were quenched with 50mM NH4Cl in PBS for 15 minutes at RT, followed by washes in PBS. Cells were then permeabilised and blocked in 3% bovine serum albumin (BSA), 0.2% Triton-X100 for 30 minutes at RT. Following this, cells were incubated for 2h at RT with primary antibody diluted in 3% BSA, 0.2% Triton X-100 in PBS. Cells were washed using 0.5% BSA, 0.05% Triton X- 100 in PBS and incubated for 1 hour at RT with fluorescently labelled secondary antibodies in 3% BSA, 0.2% Triton X-100 in PBS. Cells were then washed in 0.5% BSA, 0.05% Triton X-100 in PBS, followed by PBS and mounted using Prolong Gold (Thermo Fischer Scientific). Antibody dilutions are specified in Extended Data Table 1.

### Immunoblotting

Fully-adhered human fibroblasts were washed in PBS and immediately lysed in the tissue culture dish with 1xNUPAGE LDS sample buffer (Invitrogen) supplemented with 50mM DL-Dithiothreitol (DTT), which was then incubated at 95°C for 10 minutes.

Human sperm cells were centrifuged at 300x*g* for 15min at RT. The supernatant was carefully removed, and the pellet was resuspended in an equal volume of 1x NUPAGE LDS Sample buffer (Invitrogen) supplemented with 50 mM DTT and incubated for 10min at 95°C.

Proteins were separated using NUPAGE 4-12% Bis-Tris gels (Thermo Fischer Scientific) or 3-8% tris-acetate gels (Thermo Fischer Scientific) and MES SDS or MOPS SDS running buffer. Proteins were transferred onto PVDF membranes (BioRad) for immunoblotting and sequentially incubated in primary antibody solution, washing solution, secondary antibody solution and washing solution using an iBind Flex (Thermo Fischer Scientific) system and iBind Flex Solution kit reagents (Invitrogen). Immunoblots were imaged using a BioRad Chemidoc MP. Antibody dilutions are specified in Supp Table 1.

### Immunohistochemistry

Glass slides with pre-mounted FFPE-embedded testis sections (5μm thick) from five healthy donors with no history of infertility and between 18-55 years of age were obtained from Imperial College Healthcare Tissue Bank (Research Ethics Committee approval numbers: 22/WA/0214; ICHTB HTA license: 12275).

FFPE sections were deparaffinised, rehydrated and subjected to heat-induced antigen retrieval in 10mM Sodium Citrate, 0.05% Tween 20 (pH6). Tissue sections were then washed in 0.05% Triton-X100 in TBS for 15min with rocking at room temperature, followed by incubation in blocking solution (1%BSA, 10% Donkey Serum in TBS) for 30min at room temperature. Then they incubated with primary antibodies in 1% BSA in TBS, overnight at 4°C. Following this, tissue sections were washed in 0.05% Triton X-100 in TBS, for 10min at room temperature, with rocking, 3 times. Then they were incubated for 1h at room temperature with fluorescently labelled secondary antibodies, diluted in 1% BSA in TBS. Tissue sections were then washed in 0.05% Triton-X100 in TBS, for 10min at room temperature three times, followed by a TBS wash and diH2O. Tissue sections were stained with Hoechst 33342 (Thermo Fischer Scientific) and mounted using #1.5 coverslips and Prolong Gold Mounting media (Thermo Fischer Scientific). For antibody dilutions refer to Extended Data Table 1.

### Fluorescence Recovery After Photobleaching

Cells (both human sperm cells and human fibroblasts) were electroporated with 10kDa-FITC and 70kDa-TMR dextrans using a 4D-Nucleofector X Unit (Lonza) and reagents (Lonza). Following this, cells were let to adhere to μ-Slide 8-well high glass bottom dishes (ibidi) for 24h (human fibroblasts) or for 30min (sperm cells – ibidi surface pre-treated with PLL) at 37°C, 5%CO2. Cells were then washed in CO2- independent medium (Thermo Fischer Scientific) before imaging.

FRAP measurements were performed using a Zeiss LSM780 system, equipped with a heated stage and incubator and a 63x 1.4NA immersion oil objective. Built-in multi- band dichroic mirrors were used to reflect the excitation laser onto the sample. Fluorescent signal was collected using a multi-anode photomultiplier tube (MA-PMT) with 0.96μs pixel-dwell time.

Three frames were acquired under 488 or 561nm wavelength laser illumination before photobleaching. Selected regions of interest were exposed to full laser power, followed by 200 seconds of image acquisition. The time course of fluorescence intensity from the selected regions was recorded by Zeiss Zen 2.3 Blue software. Fluorescence intensity time traces for the regions of interest, whole-cell and background area were analysed using Zeiss Zen 2.3 Blue software (Zeiss). t1/2 values were calculated in Zen (Zeiss) by fitting a single-exponential to the normalised data.

### Stochastic Optical Reconstruction Microscopy and Cluster Analysis

Following immunofluorescence protocol, cells were imaged using a Nanoimager (ONI) system, with a 100x 1.49NA (TIRF) oil immersion objective. Imaging was performed using 488nm, 561nm and/or 640nm wavelength illumination, with acquisition of 20 000 frames and 30ms exposure time. All STORM imaging was performed using Highly Inclined and Laminated Optical Sheet (HILO) illumination with laser power 60-150mW. Imaging was performed in STORM buffer - 10% glucose, 10 mM NaCl, 50 mM Tris- HCl pH8.0, supplemented with GLOX solution (5.6% glucose oxidase and 3.4 mg/ml catalase in 50 mM NaCl, 10 mM Tris-HCl pH 8.0) and 0.1% 2-mercaptoethanol.

For two colour STORM, channel alignment was performed using a calibration procedure with TetraSpeck beads (Invitrogen). Image processing of STORM data was performed using CODI software (ONI), including drift-correction and HDBSCAN^94^ analysis. For HDBSCAN analysis of SEPT12 STORM data, a minimum cluster size was set at 15 molecules.

### Structured Illumination Microscopy

Following immunofluorescence protocol, SIM imaging was performed using a Zeiss Elyra S.1 system equipped with a 63x 1.46NA plan-apochromat oil immersion objective; 405nm, 488nm, 561nm and 647nm laser lines and a CMOS camera. 0.091μm Z-stacks were collected with 5 rotations and frame averaging of 4. Channel alignment was performed using a calibration procedure with the affine method, with mounted TetraSpeck beads (Invitrogen).

Data analysis including channel alignment, deconvolution and SR-SIM post- processing was performed in Zeiss Zen Black 2.3 (Zeiss). 3D SIM reconstructions were performed using Imaris software (Oxford Instruments).

### Confocal Microscopy

Tissue sections were imaged using an Andor Bench Top Confocal BC43 (Oxford Instruments), equipped with a 60x 1.42NA oil immersion objective, 405nm, 488nm, 561nm and 638nm lasers. For imaging acquisition Fusion software (Oxford Instruments) was used, and image analysis was performed using Imaris software (Oxford Instruments), including 3D segmentation of cells and rendering.

## Data availability

Cryo-EM map of the sperm NPC IR has been deposited in the Electron Microscopy Data Bank (EMDB) with the accession code: EMD-51638.

## Code availability

Code for converting Imod dipole picks into Warp 1.1.0 compatible star file is available here: https://zenodo.org/records/14010295

## Acknowledgments

We thank the Barford lab members for useful discussions and critical feedback. We thank Martin Beck and Bernhard Hampoelz for supporting the early stages of this project. We thank Delphine Larrieu for providing hTERT human skin fibroblasts. We thank Andrew Carter for insightful scientific discussions. We thank the cryo-EM Facility at the LMB for maintaining the electron microscopes, the Scientific Computing facility for maintaining our computer cluster and the Light Microscopy Facility for technical support with light microscopy experiments, in particular Tomás Pais de Azevedo for FRAP experiments. We acknowledge Diamond for access to the Aquilos-II of the UK national electron Bio-Imaging Centre (eBIC), in particular James B. Gilchrist for running the instrument.

This work was supported by the Medical Research Council, as part of United Kingdom Research and Innovation (also known as UK Research and Innovation) [MC_UP_1201/30; MC_UP_1201/6]. For the purpose of open access, the MRC Laboratory of Molecular Biology has applied a CC BY public copyright licence to any Author Accepted Manuscript version arising.

M.A. was funded by the UKRI Medical Research Council (MC_UP_1201/30), A.d.S. was funded by the Wellcome Trust Early Career Award (227622/Z/23/Z), D.Ba. was funded by UKRI/Medical Research Council (MC_UP_1201/6) and Cancer Research UK (C576/A14109).

Human tissue samples used in this research project were obtained from the Imperial College Healthcare Tissue Bank (ICHTB). ICHTB is supported by the National Institute for Health Research (NIHR) Biomedical Research Centre based at Imperial College

Healthcare NHS Trust and Imperial College London. ICHTB is approved by Wales REC3 to release human material for research (22/WA/0214).

## Author Contributions

M.A. and A.d.S. conceived and designed the experiments; A.d.S. and M.A. wrote the manuscript; A.d.S, M.A., T.D. and G.C. collected cryo-ET data; M.A. and A.d.S. analysed cryo-ET data; A.d.S. performed cryo-FIB milling, immunohistochemistry, STORM, SIM and FRAP experiments; O.K. performed immunoblotting, immunofluorescence and SIM experiments, supervised by A.d.S.; A.d.S performed light microscopy analysis; T.H. performed morphometrics analysis; V. L. H. performed cryo-FIB milling; A.B. and T.D. wrote scripts for tomographic analysis; P.K. supported immunoblotting work and data analysis; D.Be. supported cryo-FIB milling; D.Ba. provided guidance and technical input; M.A. supervised the study.

## Material & correspondence

Material requests and correspondence should be addressed to Matteo Allegretti.

## Ethics declarations Competing interests

The authors declare no competing interests.

Human mature sperm cells were purchased from the European Sperm Bank (https://www.europeanspermbank.com/en). The samples had been previously processed, tested by the bank according to the requirement of the national competent authorities, GMP and WHO standards, as well as EU directives: 2006/86/EC, 2006/17/EC, 2004/23/EC, 2015/565. Sperm cells were screened for morphology, genetic diseases, blood-borne viruses and motility. Consent to use the samples for research purposes was given to the European Sperm Bank.

Human testis FFPE sections from five different individuals, aged 18-55 years of age and with no history of infertility, were obtained from the Imperial College Healthcare Tissue Bank (Department of Surgery and Cancer). All the experimental procedures were approved by the Imperial College Healthcare Tissue bank (Research Ethics Committee approval numbers: 22/WA/0214; ICHTB HTA license: 12275).

## Extended Data

**Extended Data Fig. 1.**
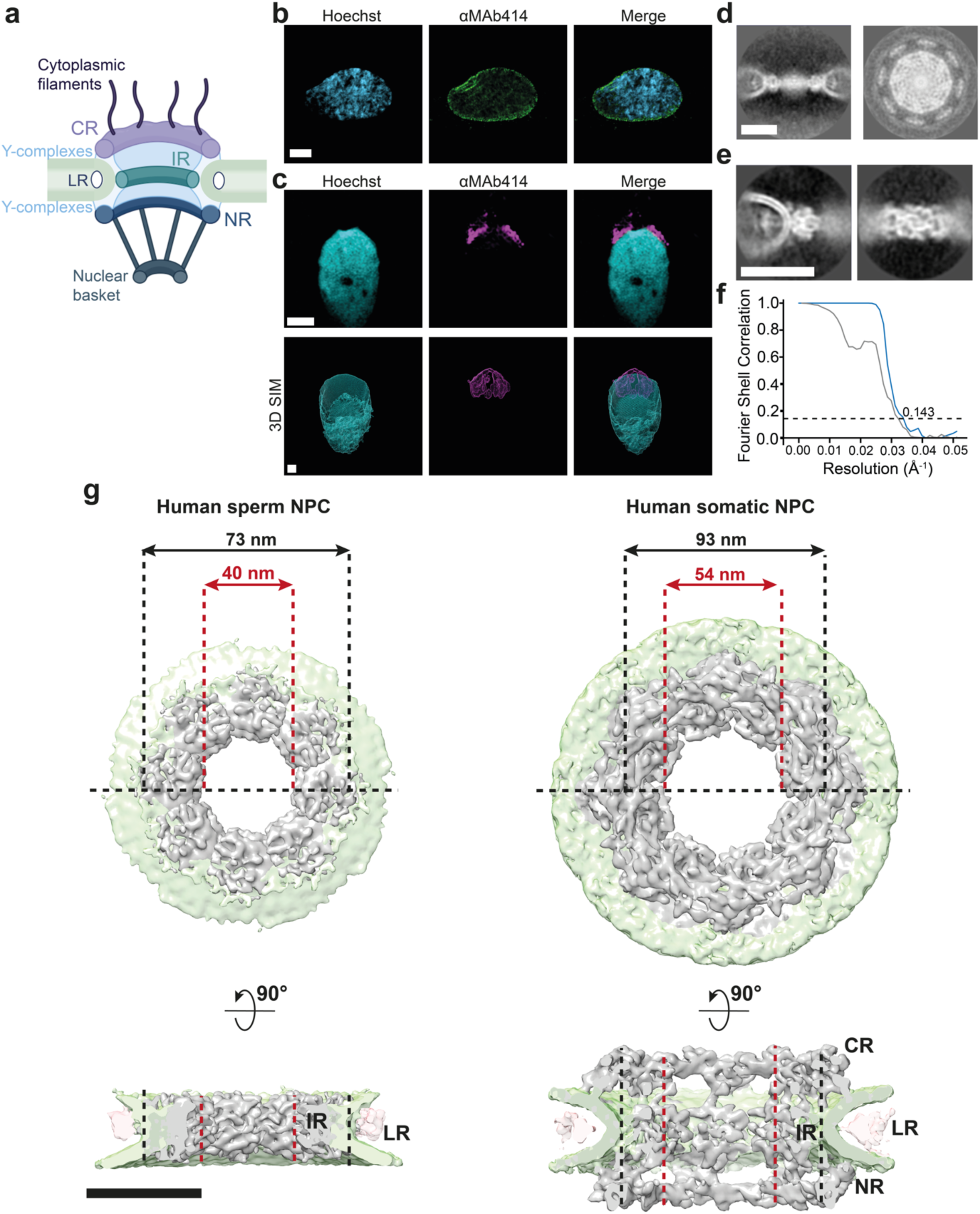
Human Sperm NPCs. a, NPC Schematic showing canonical three ring organisation: inner ring (IR), nuclear ring (NR) and cytoplasmic ring (CR). Y-complexes, cytoplasmic filaments, nuclear basket and luminal ring (LR) are also shown. **b,** Distribution of NPCs in human fibroblasts immunolabelled with MAb414 antibody (in green). DNA stained with Hoechst shown in cyan. Scale bar = 5μm. Images are representative of at least three independent experiments. **c,** Additional examples of 2D and 3D distribution of NPCs in human sperm cells using SIM in cells immunolabelled with MAb414 antibody (magenta). DNA stained with Hoechst shown in cyan. Scale bar = 1μm. Images are representative of more ten biological replicates, with at least two technical replicates. **d,** Subtomogram averaging of whole NPCs *in-situ* from human sperm cells. Scale bar = 50 nm. **e,** Sub-boxing averages of *in-situ* NPCs from human sperm cells. Scale bar = 50 nm. **f,** FSC curves for human sperm NPCs *in-situ*. FSC curve for inner ring shown in blue (∼30Å resolution) and for the luminal ring in grey (∼35Å resolution). Dotted black line displays 0.143 threshold. **g,** Comparison between the *in-situ* architecture of the human sperm NPC (this study) and the human somatic NPC structure from HEK cells (EMD-14321^13^). CR, IR and NR are indicated. Central channel diameters are shown with red dashed lines and membrane-to-membrane distance shown with black dashed lines. Scale bar = 50nm.

**Extended Data Fig. 2.**
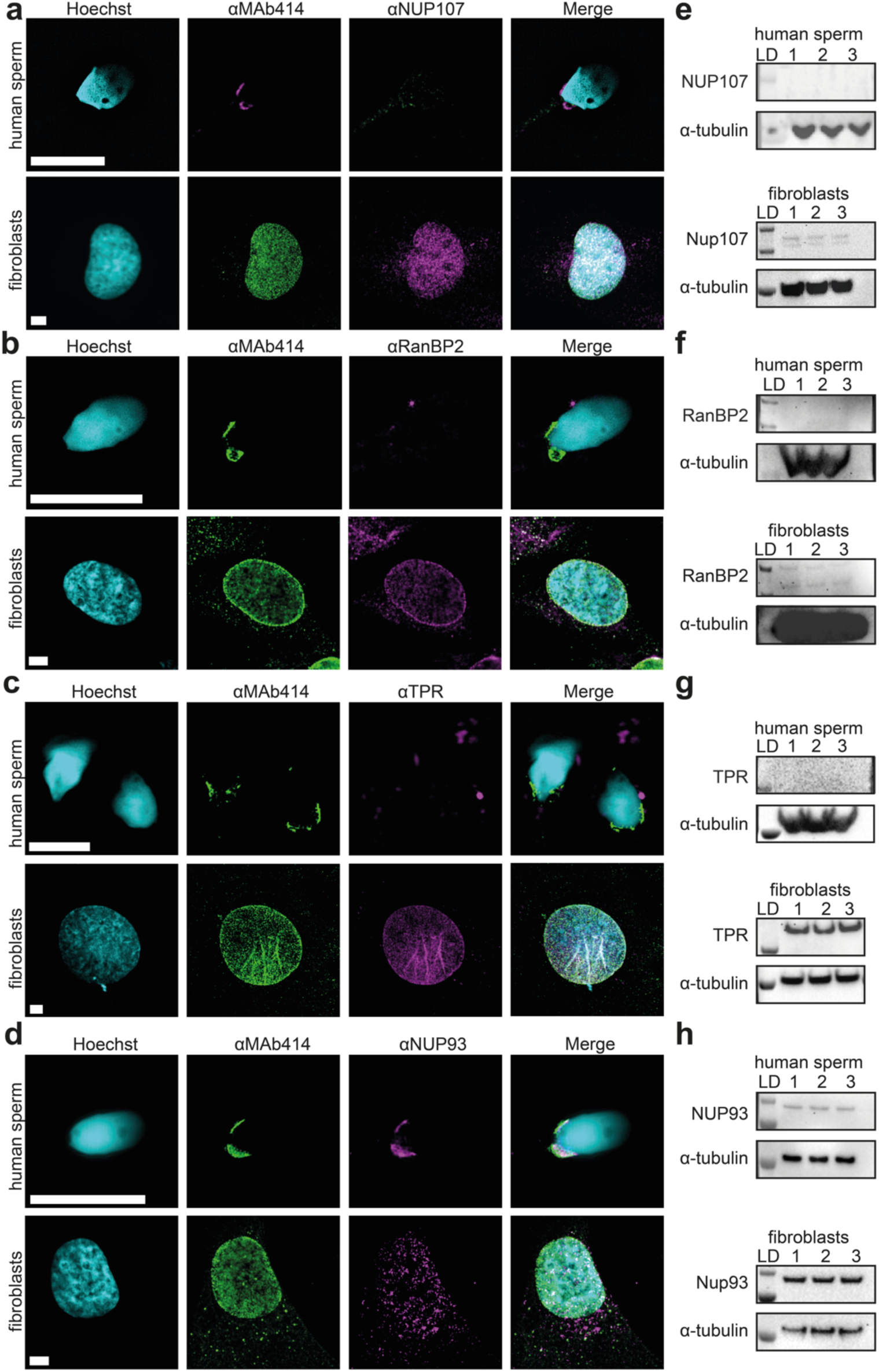
Absence of the CR and the NR in human sperm NPCs. a-d, SIM imaging of human sperm cells showing localisation of different NPC components, co-immunolabelled with MAb414 in human sperm cells. The localisation of the same NPC components in human fibroblasts is shown below. Scale bars = 5μm. Images are representative of at least three biological replicates with technical replicates (human sperm) or at least three independent experiments or technical replicates (human fibroblasts). **a,** NUP107 in human sperm and human fibroblasts; **b,** RANBP2 in human sperm and human fibroblasts; **c,** TPR in human sperm and human fibroblasts; **d,** NUP93 in human sperm and human fibroblasts. **e-h,** immunoblots for NPC components using human sperm and human fibroblasts lysate. **e,** NUP107; **f,** RanBP2; **g,** TPR; and **h,** NUP93. Replicates for three independent experiments are shown using α-tubulin as loading control.

**Extended Data Fig. 3.**
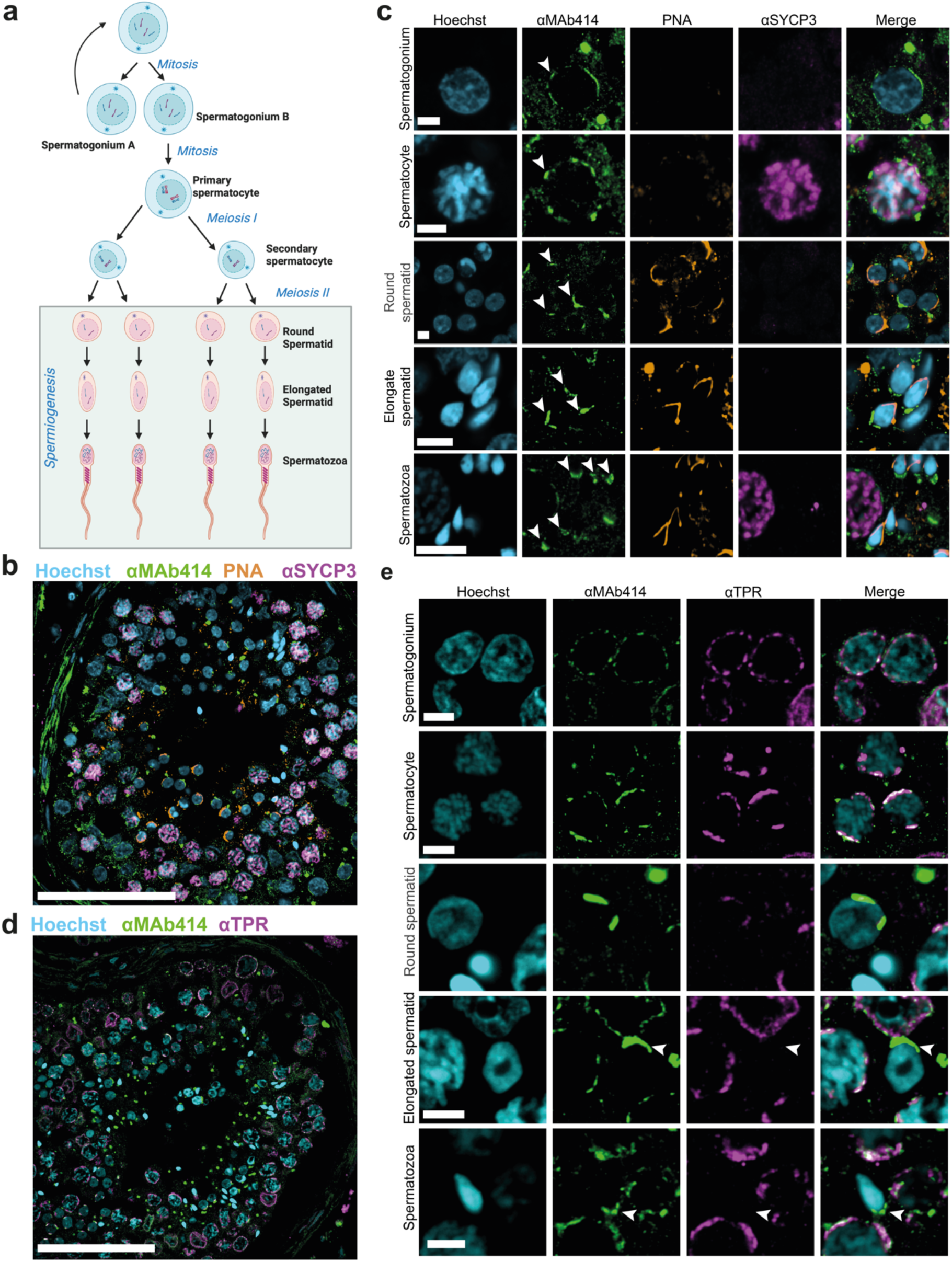
Reorganisation of NPC components during human spermatogenesis. a, Schematic of sperm cell differentiation in testis (spermatogenesis). Cell populations are indicated: spermatogonia, spermatocytes, round spermatids, elongated spermatids and spermatozoa. From round spermatid to spermatozoa stage (post-meiotic stage) the differentiation process is referred to as spermiogenesis. **b,** Confocal imaging of a representative seminiferous tubule from human testis sections, showing NPCs in green (MAb414), DNA stained with Hoechst shown in cyan, meiotic marker SYCP3 and Peanut Agglutinin (PNA) as an acrosomal marker (orange). Scale bar = 50μm. Images are representative of at least three independent experiments. **c,** Detail of subpopulations of cells labelled as described in panel *a*, showing reorganisation of NPCs (in green, arrowheads) relative to the acrosome (orange). Hoechst staining of DNA is shown in cyan. Scale bar = 50μm. **d,** Immunohistochemistry and confocal imaging of a representative human seminiferous tubule. NPCs (MAb414) are shown in green and nuclear basket component TPR is shown in magenta. Scale bar = 50μm. Images are representative of seminiferous tubules from at least two human donors, with technical replicates. **e,** Detail of subpopulations of cells in human testes, labelled as described in **(c)**. Arrowheads indicate loss of co-localisation between MAb414 and TPR signals. DNA is shown in cyan (Hoechst). Scale bar = 5μm.

**Extended Data Fig. 4.**
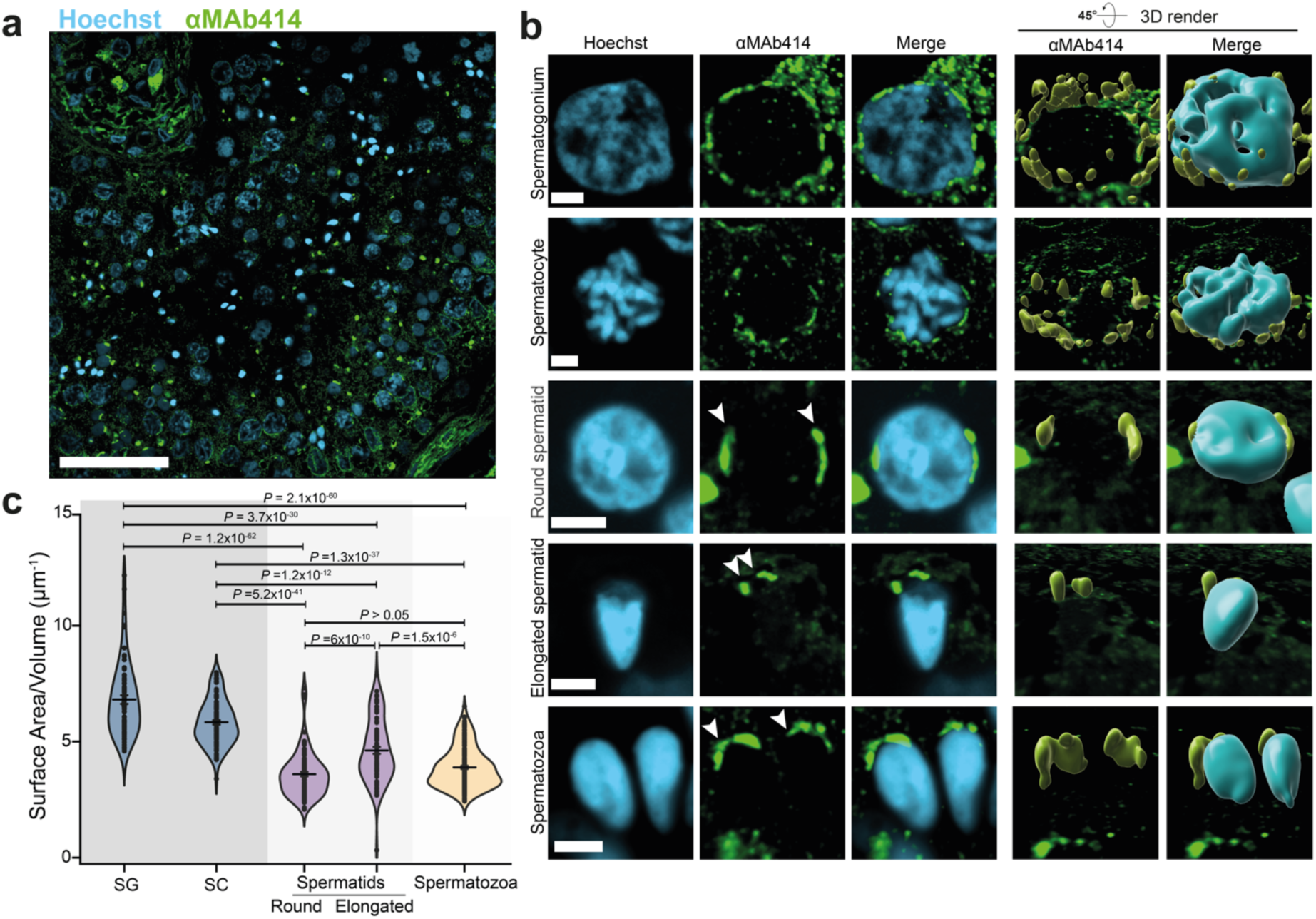
Re-organisation and clustering of NPCs during human spermatogenesis. a, Confocal imaging of a representative seminiferous tubule in testis tissue from healthy human donors. DNA staining with Hoechst is shown in cyan and NPCs are immunolabelled with MAb414 antibody (green). Scale bar = 50μm. Data are representative of seminiferous tubules from five human donors, with at least three technical replicates. **b,** Detail of different cell populations within seminiferous tubules of human testicular tissue, showing distribution of NPCs at the NE. 3D rendering of NPC regions and DNA also shown (right panels). Reorganisation of NPCs in post-mitotic cells (arrowheads - round spermatids to spermatozoa) can be observed. Scale bars = 5μm. **c,** Quantification of surface area/volume ratio of NPC regions for each cell subpopulation in human testicular tissue – spermatogonia (SG), spermatocytes (SC), round and elongated spermatids and spermatozoa. Mean ± SE values: spermatogonia (SG) = 6.81 ± 0.19 μm^-1^, n=72; spermatocytes (SC) = 5.83 ± 0.10 μm^-1^, n=100; round spermatids = 3.60 ± 0.09 μm^-1^, n=112; elongated spermatids = 4.62 ± 0.15 μm^-1^, n=80; spermatozoa = 3.89 ± 0.67 μm^-1^, n=169. Data used for plotting was collected from five biological replicates, with at least three technical replicates. Statistical Analysis: ANOVA Tukey test; *P*-values shown.

**Extended Data Fig. 5.**
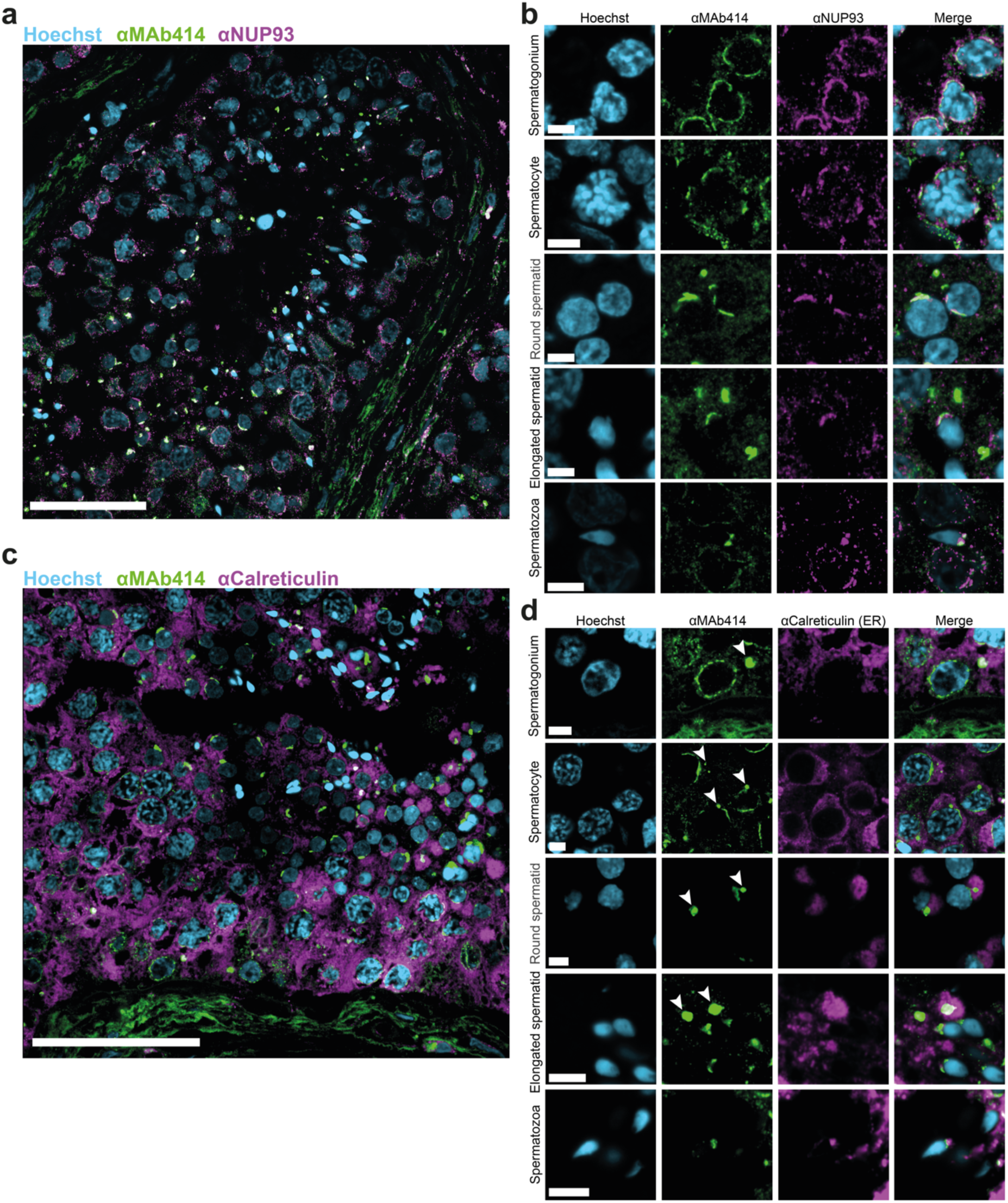
Localisation of NPC components and ER during spermatogenesis. a, Immunohistochemistry and confocal imaging of representative human seminiferous tubule in testes sections, showing localisation of NPCs (MAb414, green) and IR component NUP93 (magenta). Hoechst is shown in cyan. Scale bar = 50μm. Data are representative of seminiferous tubules from at least three human donors. **b,** Detail of cellular sub-populations in human testis, labelled as described in **(a)** showing co-localisation of MAb414 and NUP93 at all steps of differentiation. **c,** Overview of a representative human seminiferous tubule showing endoplasmic reticulum (ER) marker calreticulin in magenta and MAb414 (green). Scale bar = 50μm. Data are representative of three independent experiments. **d,** Detail of differentiating cells in human testis with labelling as described in **(c)**. Arrowheads indicate co-localisation of MAb414 signal (green) and calreticulin (magenta) in the cytoplasm of all cell types, except mature spermatozoa. Scale bar = 5μm.

**Extended Data Fig. 6.**
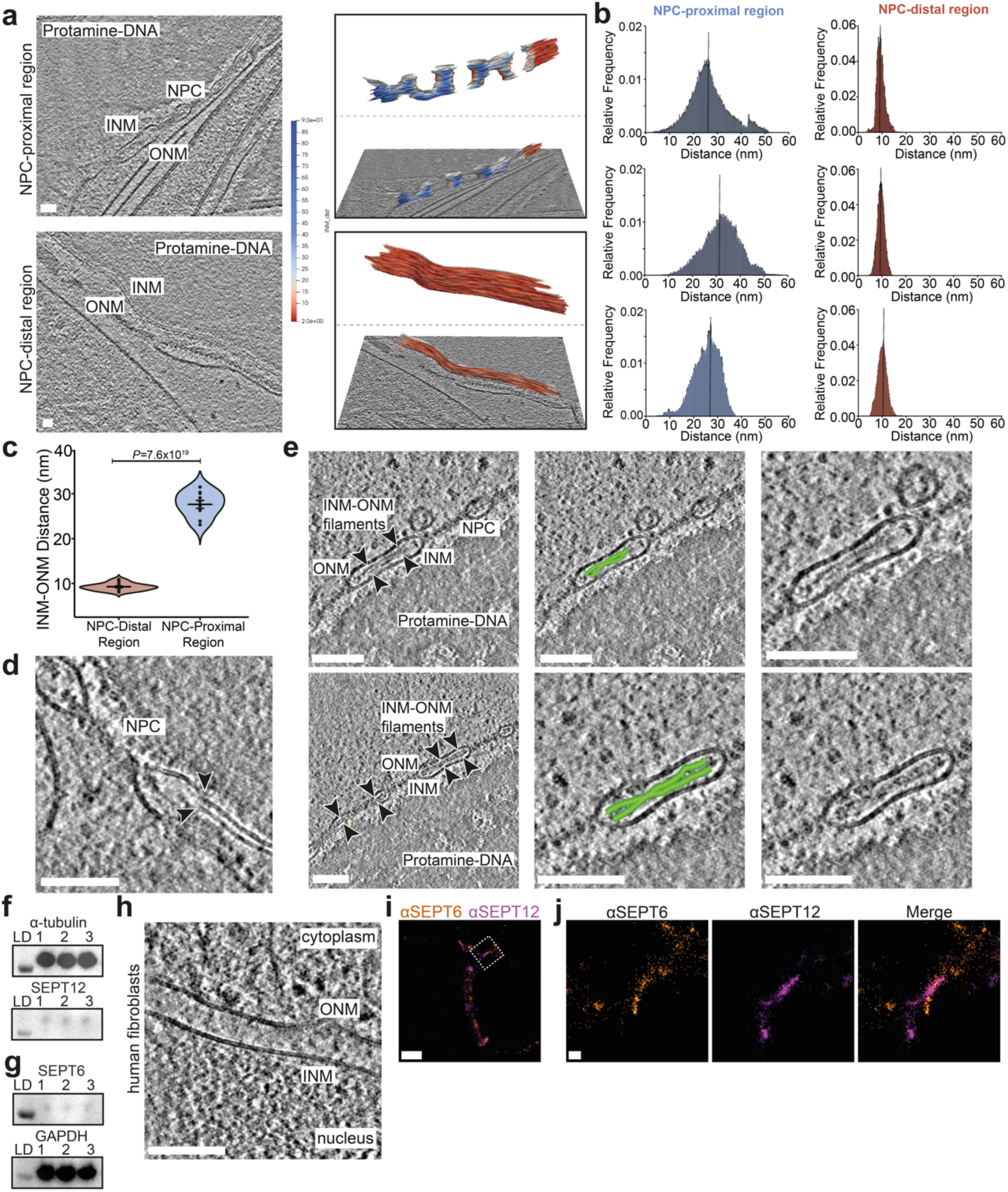
INM-ONM intermembrane filaments. a, Tomograms showing INM and ONM spacing examples relative to their proximity to NPCs. NPC-distal and proximal regions considered as >200nm and <150nm distance, respectively, from the centre of any NPC. Morphometrics segmentation and heat map of INM-ONM distances shown on the right panel. Blue colour indicates wider INM-ONM distances; brown indicates shorter distances. Scale bars = 100nm. **b,** Example histograms, plotted from morphometrics analysis of tomographic data, showing distribution of INM-ONM distances. Each histogram represents the distributions of INM-ONM measurements for a single tomogram. Histograms for NPC-distal regions shown in brown, and for NPC-proximal regions in blue. Modal values are shown as black dashed lines. **c,** Violin plots with distribution of modal values obtained from histograms, following morphometrics analysis of tomographic data. Mean ± SE values: NPC-distal region = 9.24 ± 0.20nm, n = 16 cells; NPC-proximal region = 27.64 ± 2.76nm, n = 10 cells. Data from at least three biological replicates (donors). Statistical Analysis: t- test; *P-*values are shown. **d,** Tomogram detail of transition region (arrowheads) between narrow INM-ONM and wider INM-ONM regions. Scale bar = 100nm. **e,** Tomography data of NPC regions in human sperm cells showing the presence of filamentous structures within the INM-ONM intermembrane space (arrowheads). Mid-panel shows tomogram segmentation of filaments, and a zoomed-in image is shown on the right-panel. Scale bars = 100nm. **f,** Immunoblot for SEPT12 using human sperm lysate. Three replicates are shown with α-tubulin as a loading control. **g,** Immunoblot for SEPT6 using human sperm lysate. Three replicates are shown with GAPDH as a loading control. **h,** Tomographic slice of human fibroblast showing NE region. Image shows a representative slice from more than twenty tomograms. Scale bar = 100 nm. **g,** STORM imaging of a human sperm cell showing localisation of SEPT12 (magenta) and SEPT6 (orange). Dashed white-line rectangle shows detail of an NPC region. Scale bar = 1μm. Data are representative of at least three independent experiments. **h,** STORM image showing localisation of SEPT12 (magenta) relative to SEPT6 (orange) in NPC region selected from **(h)** – dashed-white rectangle. Scale bar = 100nm.

**Extended Data Fig. 7.**
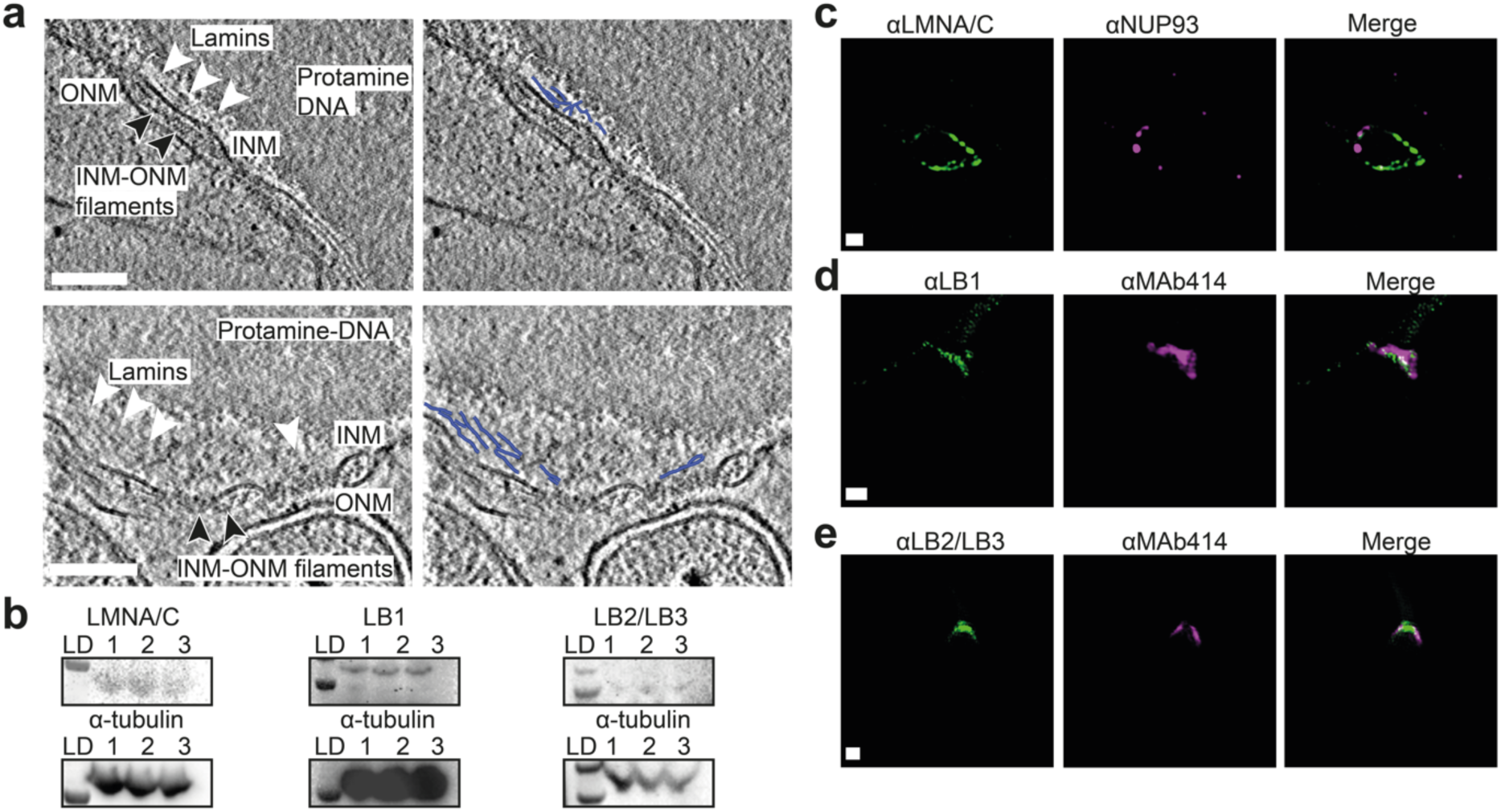
Lamins in human sperm cells. a, Observation of nuclear lamina (white arrowheads) at the NE of human sperm cells in tomographic data. INM-ONM intermembrane filaments are indicated with black arrowheads. Right panels show segmentation of lamina filaments (blue lines). Scale bars = 100 nm. **b,** immunoblots against nuclear lamina components Lamin A/C (LMNA/C), Lamin B1 (LB1), and Lamin B2/B3 (LB2/B3) using human sperm lysate, with three replicates. α-tubulin shown as loading control. **c-e,** SIM imaging of immunofluorescently labelled human sperm cells showing localisation of nuclear lamina components (green) **c,** LMAN/C **d,** LB1 **e,** LB2/LB3 relative to NPCs (MAb414 or NUP93, magenta). Scale bars = 1μm. Images are representative of at least three independent experiments.

**Extended Data Fig. 8.**
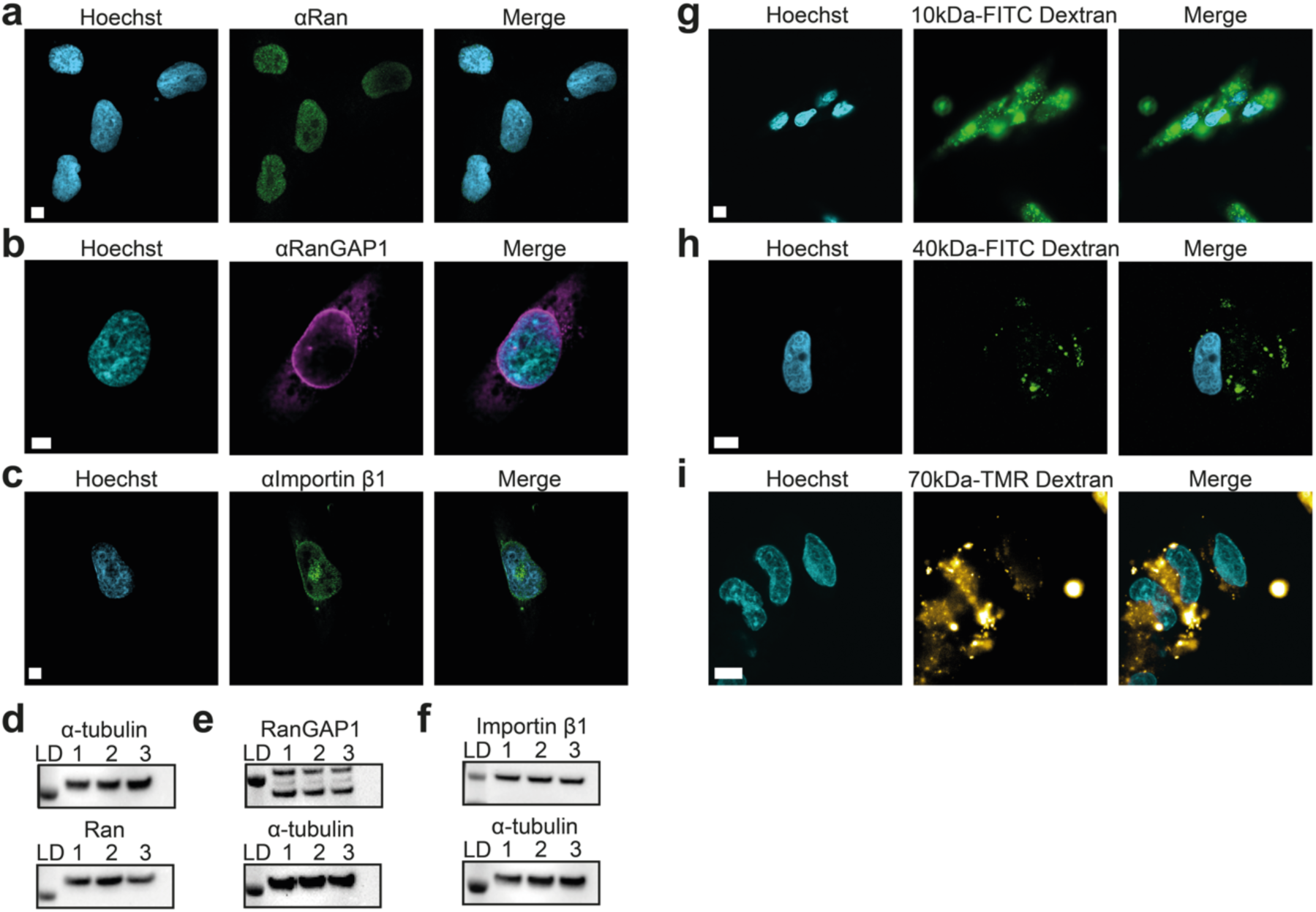
Active transport and diffusion in human fibroblasts. a-c, Immunofluorescence and confocal imaging of different nuclear transport components: **a,** Ran showing nuclear localisation during interphase in human fibroblasts; **b,** RanGAP1 showing localisation at the NE and low cytoplasm signal; **c,** Importin β1, showing nuclear signal. Scale bars = 5μm. Images are representative of at least three independent experiments (technical replicates). **d-f,** Western Blots against **d,** Ran; **e,** RanGAP1; and **f,** Importin β1 using human fibroblasts lysate, shown with three replicates and α-tubulin as loading control. **g-i,** Human fibroblasts treated with digitonin for the delivery of **g,** 10kDa-FITC; **h,** 40kDa-FITC and **i,** 70kDa- TMR dextrans. Signal for 10kDa-FITC Dextran, but not 40kDa-FITC or 70kDa-TMR Dextran, can be observed in the nucleus. Scale bars =5μm.

**Extended Data Table 1.** Overview of antibodies used in this work.

| Antibody | Catalogue# | Company | Dilutions |  |  |
| --- | --- | --- | --- | --- | --- |
|  |  |  | Immunofluorescence | Immunoblotting | Immunohistochemistry |
| mouse MAb414 | ab24609 | Abcam | 1:200 | - | 1:200 |
| rabbit anti-NUP107 | DF4250 | Affinity Biosciences | 1:100 | 1:500 | 1:100 |
| rabbit anti-NUP93 | PA553674 | Invitrogen | 1:200 | 1:200 | 1:200 |
| rabbit anti-NUP358 (RanBP2) | HPA 067564 | Sigma | 1:50 | 1:500 | 1:100 |
| rabbit anti-TPR | ab84516 | Abcam | 1:50 | 1:500 | 1:100 |
| rabbit anti-SEPT12 | PA5-31504 | Invitrogen | 1:200 | 1:200 | - |
| mouse conjugated anti-SEPT6-AF594 | sc-514781 | Santa Cruz | 1:100 | - | - |
| rabbit anti-SEPT6 | PA5-52177 | Invitrogen | - | 1:2000 | - |
| rabbit anti-LB1 | ab16048 | Abcam | 1:100 | 1:1000 | - |
| rabbit anti-LB2/LB3 | ab151735 | Abcam | 1:100 | 1:1000 | - |
| mouse anti-LMNA/C | sc-7292 | Santa Cruz | 1:100 | 1:1000 | - |
| mouse anti-Ran | sc-271376 | Santa Cruz | 1:100 | 1:1000 | - |
| mouse anti-Importin β1 | MA3-070 | Invitrogen | 1:100 | 1:1000 | - |
| rabbit anti-RanGAP1 | MA5-34762 | ThermoFisher | 1:50 | 1:500 | - |
| mouse anti-GAPDH | MA4300 | Invitrogen | - | 1:5000 | - |
| mouse anti-alpha tubulin | A11126 | Invitrogen | - | 1:5000 | - |
| rabbit anti-alpha tubulin | ab52866 | Abcam | - | 1:5000 | - |
| mouse anti-FLAG | 66008-4 | Proteintech | 1:200 | - | - |
| rabbit anti-FLAG | 20543-1 | Proteintech | 1:200 | - | - |
| goat anti-mouse HRP | ab97023 | Abcam | - | 1:10000 | - |
| goat anti-rabbit HRP | ab6721 | Abcam | - | 1:15000 | - |
| donkey anti-mouse Alexa 488 | ab181289 | Abcam | 1:500 | - | 1:500 |
| donkey anti-mouse Alexa 555 | ab150110 | Abcam | 1:500 | - | 1:500 |
| donkey anti-mouse Alexa 647 | ab150111 | Abcam | 1:500 | - | 1:500 |
| donkey anti-rabbit Alexa 488 | ab181346 | Abcam | 1:500 | - | 1:500 |
| donkey anti-rabbit Alexa 555 | ab150074 | Abcam | 1:500 | - | 1:500 |
| donkey anti-rabbit Alexa 647 | ab181347 | Abcam | 1:500 | - | 1:500 |
| donkey anti-mouse Atto 488 | 62197 | Abcam | 1:500 | - | - |
| donkey anti-rabbit Atto-488 | 18772 | Abcam | 1:500 | - | - |

## REFERENCES

1. Wente, S.R., and Rout, M.P. (2010). The nuclear pore complex and nuclear transport. Cold Spring Harb Perspect Biol 2, a000562. 10.1101/cshperspect.a000562.

2. Mosalaganti, S., Obarska-Kosinska, A., Siggel, M., Taniguchi, R., Turonova, B., Zimmerli, C.E., Buczak, K., Schmidt, F.H., Margiotta, E., Mackmull, M.T., et al. (2022). AI-based structure prediction empowers integrative structural analysis of human nuclear pores. Science 376, eabm9506. 10.1126/science.abm9506.

3. Huang, G., Zeng, C., and Shi, Y. (2023). Structure of the nuclear pore complex goes atomic. Curr Opin Struct Biol 78, 102523. 10.1016/j.sbi.2022.102523.

4. Okazaki, R., Yamazoe, K., and Inoue, Y.H. (2020). Nuclear Export of Cyclin B Mediated by the Nup62 Complex Is Required for Meiotic Initiation in Drosophila Males. Cells 9. 10.3390/cells9020270.

5. Yang, W.X., Jefferson, H., and Sperry, A.O. (2006). The molecular motor KIFC1 associates with a complex containing nucleoporin NUP62 that is regulated during development and by the small GTPase RAN. Biol Reprod 74, 684–690. 10.1095/biolreprod.105.049312.

6. Lin, D.H., and Hoelz, A. (2019). The Structure of the Nuclear Pore Complex (An Update). Annu Rev Biochem 88, 725–783. 10.1146/annurev-biochem-062917-011901.

7. Raices, M., and D’Angelo, M.A. (2017). Nuclear pore complexes and regulation of gene expression. Curr Opin Cell Biol 46, 26–32. 10.1016/j.ceb.2016.12.006.

8. Hampoelz, B., Andres-Pons, A., Kastritis, P., and Beck, M. (2019). Structure and Assembly of the Nuclear Pore Complex. Annu Rev Biophys 48, 515–536. 10.1146/annurev-biophys-052118-115308.

9. Zhang, Y., Li, S., Zeng, C., Huang, G., Zhu, X., Wang, Q., Wang, K., Zhou, Q., Yan, C., Zhang, W., et al. (2020). Molecular architecture of the luminal ring of the Xenopus laevis nuclear pore complex. Cell Res 30, 532–540. 10.1038/s41422-020-0320-y.

10. 10. Bui, K.H., von Appen, A., DiGuilio, A.L., Ori, A., Sparks, L., Mackmull, M.T., Bock, T., Hagen, W., Andres-Pons, A., Glavy, J.S., and Beck, M. (2013). Integrated structural analysis of the human nuclear pore complex scaffold. Cell 155, 1233–1243. 10.1016/j.cell.2013.10.055.

11. Bley, C.J., Nie, S., Mobbs, G.W., Petrovic, S., Gres, A.T., Liu, X., Mukherjee, S., Harvey, S., Huber, F.M., Lin, D.H., et al. (2022). Architecture of the cytoplasmic face of the nuclear pore. Science 376, eabm9129. 10.1126/science.abm9129.

12. Fontana, P., Dong, Y., Pi, X., Tong, A.B., Hecksel, C.W., Wang, L., Fu, T.M., Bustamante, C., and Wu, H. (2022). Structure of cytoplasmic ring of nuclear pore complex by integrative cryo-EM and AlphaFold. Science 376, eabm9326. 10.1126/science.abm9326.

13. Zhu, X., Huang, G., Zeng, C., Zhan, X., Liang, K., Xu, Q., Zhao, Y., Wang, P., Wang, Q., Zhou, Q., et al. (2022). Structure of the cytoplasmic ring of the Xenopus laevis nuclear pore complex. Science 376, eabl8280. 10.1126/science.abl8280.

14. Otsuka, S., Bui, K.H., Schorb, M., Hossain, M.J., Politi, A.Z., Koch, B., Eltsov, M., Beck, M., and Ellenberg, J. (2016). Nuclear pore assembly proceeds by an inside-out extrusion of the nuclear envelope. Elife 5. 10.7554/eLife.19071.

15. Otsuka, S., Tempkin, J.O.B., Zhang, W., Politi, A.Z., Rybina, A., Hossain, M.J., Kueblbeck, M., Callegari, A., Koch, B., Morero, N.R., et al. (2023). A quantitative map of nuclear pore assembly reveals two distinct mechanisms. Nature 613, 575–581. 10.1038/s41586-022-05528-w.

16. 16. Singh, D., Soni, N., Hutchings, J., Echeverria, I., Shaikh, F., Duquette, M., Suslov, S., Li, Z., van Eeuwen, T., Molloy, K., et al. (2024). The molecular architecture of the nuclear basket. Cell. 10.1016/j.cell.2024.07.020.

17. Li, Y., Aksenova, V., Tingey, M., Yu, J., Ma, P., Arnaoutov, A., Chen, S., Dasso, M., and Yang, W. (2021). Distinct roles of nuclear basket proteins in directing the passage of mRNA through the nuclear pore. Proc Natl Acad Sci U S A 118. 10.1073/pnas.2015621118.

18. Walther, T.C., Fornerod, M., Pickersgill, H., Goldberg, M., Allen, T.D., and Mattaj, I.W. (2001). The nucleoporin Nup153 is required for nuclear pore basket formation, nuclear pore complex anchoring and import of a subset of nuclear proteins. EMBO J 20, 5703–5714. 10.1093/emboj/20.20.5703.

19. Terry, L.J., Shows, E.B., and Wente, S.R. (2007). Crossing the nuclear envelope: hierarchical regulation of nucleocytoplasmic transport. Science 318, 1412–1416. 10.1126/science.1142204.

20. 20. von Appen, A., Kosinski, J., Sparks, L., Ori, A., DiGuilio, A.L., Vollmer, B., Mackmull, M.T., Banterle, N., Parca, L., Kastritis, P., et al. (2015). In situ structural analysis of the human nuclear pore complex. Nature 526, 140–143. 10.1038/nature15381.

21. Hutten, S., Flotho, A., Melchior, F., and Kehlenbach, R.H. (2008). The Nup358- RanGAP complex is required for efficient importin alpha/beta-dependent nuclear import. Mol Biol Cell 19, 2300–2310. 10.1091/mbc.e07-12-1279.

22. Hampoelz, B., Mackmull, M.T., Machado, P., Ronchi, P., Bui, K.H., Schieber, N., Santarella-Mellwig, R., Necakov, A., Andres-Pons, A., Philippe, J.M., et al. (2016). Pre-assembled Nuclear Pores Insert into the Nuclear Envelope during Early Development. Cell 166, 664–678. 10.1016/j.cell.2016.06.015.

23. Hampoelz, B., Schwarz, A., Ronchi, P., Bragulat-Teixidor, H., Tischer, C., Gaspar, I., Ephrussi, A., Schwab, Y., and Beck, M. (2019). Nuclear Pores Assemble from Nucleoporin Condensates During Oogenesis. Cell 179, 671–686 e617. 10.1016/j.cell.2019.09.022.

24. Sachweh, J., Börmel, M., Klumpe, S., Becker, A., Taniguchi, R., Kubańska, M.A., Pintschovius, V., Kaindl, E., Plitzko, J.M., Wilfling, F., et al. (2024). The small GTPase Ran defines Nuclear Pore Complex Asymmetry. bioRxiv, 2024.2010.2010.617378. 10.1101/2024.10.10.617378.

25. Ren, H., Xin, G., Jia, M., Zhu, S., Lin, Q., Wang, X., Jiang, Q., and Zhang, C. (2019). Postmitotic annulate lamellae assembly contributes to nuclear envelope reconstitution in daughter cells. J Biol Chem 294, 10383–10391. 10.1074/jbc.AC119.008171.

26. Mahamid, J., Pfeffer, S., Schaffer, M., Villa, E., Danev, R., Cuellar, L.K., Forster, F., Hyman, A.A., Plitzko, J.M., and Baumeister, W. (2016). Visualizing the molecular sociology at the HeLa cell nuclear periphery. Science 351, 969–972. 10.1126/science.aad8857.

27. Mosalaganti, S., Kosinski, J., Albert, S., Schaffer, M., Strenkert, D., Salome, P.A., Merchant, S.S., Plitzko, J.M., Baumeister, W., Engel, B.D., and Beck, M. (2018). In situ architecture of the algal nuclear pore complex. Nat Commun 9, 2361. 10.1038/s41467- 018-04739-y.

28. Akey, C.W., Singh, D., Ouch, C., Echeverria, I., Nudelman, I., Varberg, J.M., Yu, Z., Fang, F., Shi, Y., Wang, J., et al. (2022). Comprehensive structure and functional adaptations of the yeast nuclear pore complex. Cell 185, 361–378 e325. 10.1016/j.cell.2021.12.015.

29. Allegretti, M., Zimmerli, C.E., Rantos, V., Wilfling, F., Ronchi, P., Fung, H.K.H., Lee, C.W., Hagen, W., Turonova, B., Karius, K., et al. (2020). In-cell architecture of the nuclear pore and snapshots of its turnover. Nature 586, 796–800. 10.1038/s41586-020-2670-5.

30. Schuller, A.P., Wojtynek, M., Mankus, D., Tatli, M., Kronenberg-Tenga, R., Regmi, S.G., Dip, P.V., Lytton-Jean, A.K.R., Brignole, E.J., Dasso, M., et al. (2021). The cellular environment shapes the nuclear pore complex architecture. Nature 598, 667–671. 10.1038/s41586-021-03985-3.

31. Zimmerli, C.E., Allegretti, M., Rantos, V., Goetz, S.K., Obarska-Kosinska, A., Zagoriy, I., Halavatyi, A., Hummer, G., Mahamid, J., Kosinski, J., and Beck, M. (2021). Nuclear pores dilate and constrict in cellulo. Science 374, eabd9776. 10.1126/science.abd9776.

32. 32. Elosegui-Artola, A., Andreu, I., Beedle, A.E.M., Lezamiz, A., Uroz, M., Kosmalska, A.J., Oria, R., Kechagia, J.Z., Rico-Lastres, P., Le Roux, A.L., et al. (2017). Force Triggers YAP Nuclear Entry by Regulating Transport across Nuclear Pores. Cell 171, 1397–1410 e1314. 10.1016/j.cell.2017.10.008.

33. 33. Kreysing, J.P., Heidari, M., Zila, V., Cruz-Leon, S., Obarska-Kosinska, A., Laketa, V., Welsch, S., Köfinger, J., Turoňová, B., Hummer, G., et al. (2024). Passage of the HIV capsid cracks the nuclear pore. bioRxiv, 2024.2004.2023.590733. 10.1101/2024.04.23.590733.

34. Shen, W., Gong, B., Xing, C., Zhang, L., Sun, J., Chen, Y., Yang, C., Yan, L., Chen, L., Yao, L., et al. (2022). Comprehensive maturity of nuclear pore complexes regulates zygotic genome activation. Cell 185, 4954–4970.e4920. 10.1016/j.cell.2022.11.011.

35. 35. Taniguchi, R., Orniacki, C., Kreysing, J.P., Zila, V., Zimmerli, C.E., Böhm, S., Turoňová, B., Kräusslich, H.-G., Doye, V., and Beck, M. (2024). Nuclear pores safeguard the integrity of the nuclear envelope. bioRxiv, 2024.2002.2005.578890. 10.1101/2024.02.05.578890.

36. 36. Morgan, K.J., Carley, E., Coyne, A.N., Rothstein, J.D., Lusk, C.P., and King, M.C. (2024). Visualizing nuclear pore complex plasticity with Pan-Expansion Microscopy. bioRxiv, 2024.2009.2018.613744. 10.1101/2024.09.18.613744.

37. O’Donnell, L. (2014). Mechanisms of spermiogenesis and spermiation and how they are disturbed. Spermatogenesis 4, e979623. 10.4161/21565562.2014.979623.

38. 38. Pereira, C.D., Serrano, J.B., Martins, F., da Cruz, E.S.O.A.B., and Rebelo, S. (2019). Nuclear envelope dynamics during mammalian spermatogenesis: new insights on male fertility. Biol Rev Camb Philos Soc 94, 1195–1219. 10.1111/brv.12498.

39. Morgan, M., Kumar, L., Li, Y., and Baptissart, M. (2021). Post-transcriptional regulation in spermatogenesis: all RNA pathways lead to healthy sperm. Cell Mol Life Sci 78, 8049–8071. 10.1007/s00018-021-04012-4.

40. Whiley, P.A., Miyamoto, Y., McLachlan, R.I., Jans, D.A., and Loveland, K.L. (2012). Changing subcellular localization of nuclear transport factors during human spermatogenesis. Int J Androl 35, 158–169. 10.1111/j.1365-2605.2011.01202.x.

41. Ho, H.C. (2010). Redistribution of nuclear pores during formation of the redundant nuclear envelope in mouse spermatids. J Anat 216, 525–532. 10.1111/j.1469-7580.2009.01204.x.

42. Arafah, K., Lopez, F., Cazin, C., Kherraf, Z.E., Tassistro, V., Loundou, A., Arnoult, C., Thierry-Mieg, N., Bulet, P., Guichaoua, M.R., and Ray, P.F. (2021). Defect in the nuclear pore membrane glycoprotein 210-like gene is associated with extreme uncondensed sperm nuclear chromatin and male infertility: a case report. Hum Reprod 36, 693–701. 10.1093/humrep/deaa329.

43. Miyamoto, Y., Boag, P.R., Hime, G.R., and Loveland, K.L. (2012). Regulated nucleocytoplasmic transport during gametogenesis. Biochim Biophys Acta 1819, 616–630. 10.1016/j.bbagrm.2012.01.015.

44. Cavazza, T., Takeda, Y., Politi, A.Z., Aushev, M., Aldag, P., Baker, C., Choudhary, M., Bucevičius, J., Lukinavičius, G., Elder, K., et al. (2021). Parental genome unification is highly error-prone in mammalian embryos. Cell 184, 2860–2877.e2822. 10.1016/j.cell.2021.04.013.

45. Dultz, E., and Ellenberg, J. (2010). Live imaging of single nuclear pores reveals unique assembly kinetics and mechanism in interphase. J Cell Biol 191, 15–22. 10.1083/jcb.201007076.

46. Zila, V., Margiotta, E., Turonova, B., Muller, T.G., Zimmerli, C.E., Mattei, S., Allegretti, M., Borner, K., Rada, J., Muller, B., et al. (2021). Cone-shaped HIV-1 capsids are transported through intact nuclear pores. Cell 184, 1032–1046 e1018. 10.1016/j.cell.2021.01.025.

47. Chemes, H.E., Fawcett, D.W., and Dym, M. (1978). Unusual features of the nuclear envelope in human spermatogenic cells. Anat Rec 192, 493–512. 10.1002/ar.1091920404.

48. Barad, B.A., Medina, M., Fuentes, D., Wiseman, R.L., and Grotjahn, D.A. (2023). Quantifying organellar ultrastructure in cryo-electron tomography using a surface morphometrics pipeline. J Cell Biol 222. 10.1083/jcb.202204093.

49. Turgay, Y., Eibauer, M., Goldman, A.E., Shimi, T., Khayat, M., Ben-Harush, K., Dubrovsky-Gaupp, A., Sapra, K.T., Goldman, R.D., and Medalia, O. (2017). The molecular architecture of lamins in somatic cells. Nature 543, 261–264. 10.1038/nature21382.

50. Lin, Y.-H., Kuo, Y.-C., Chiang, H.-S., and Kuo, P.-L. (2011). The role of the septin family in spermiogenesis. Spermatogenesis 1, 298–302. 10.4161/spmg.1.4.18326.

51. Chen, H., Li, P., Du, X., Zhao, Y., Wang, L., Tian, Y., Song, X., Shuai, L., Bai, X., and Chen, L. (2022). Homozygous Loss of Septin12, but not its Haploinsufficiency, Leads to Male Infertility and Fertilization Failure. Front Cell Dev Biol 10, 850052. 10.3389/fcell.2022.850052.

52. Lin, Y.H., Chou, C.K., Hung, Y.C., Yu, I.S., Pan, H.A., Lin, S.W., and Kuo, P.L. (2011). SEPT12 deficiency causes sperm nucleus damage and developmental arrest of preimplantation embryos. Fertil Steril 95, 363–365. 10.1016/j.fertnstert.2010.07.1064.

53. Shen, Y.R., Wang, H.Y., Kuo, Y.C., Shih, S.C., Hsu, C.H., Chen, Y.R., Wu, S.R., Wang, C.Y., and Kuo, P.L. (2017). SEPT12 phosphorylation results in loss of the septin ring/sperm annulus, defective sperm motility and poor male fertility. PLoS Genet 13, e1006631. 10.1371/journal.pgen.1006631.

54. Yeh, C.H., Kuo, P.L., Wang, Y.Y., Wu, Y.Y., Chen, M.F., Lin, D.Y., Lai, T.H., Chiang, H.S., and Lin, Y.H. (2015). SEPT12/SPAG4/LAMINB1 complexes are required for maintaining the integrity of the nuclear envelope in postmeiotic male germ cells. PLoS One 10, e0120722. 10.1371/journal.pone.0120722.

55. Lai, T.H., Wu, Y.Y., Wang, Y.Y., Chen, M.F., Wang, P., Chen, T.M., Wu, Y.N., Chiang, H.S., Kuo, P.L., and Lin, Y.H. (2016). SEPT12-NDC1 Complexes Are Required for Mammalian Spermiogenesis. Int J Mol Sci 17. 10.3390/ijms17111911.

56. Sirajuddin, M., Farkasovsky, M., Hauer, F., Kuhlmann, D., Macara, I.G., Weyand, M., Stark, H., and Wittinghofer, A. (2007). Structural insight into filament formation by mammalian septins. Nature 449, 311–315. 10.1038/nature06052.

57. Tanaka-Takiguchi, Y., Kinoshita, M., and Takiguchi, K. (2009). Septin-mediated uniform bracing of phospholipid membranes. Curr Biol 19, 140–145. 10.1016/j.cub.2008.12.030.

58. Kuo, Y.C., Lin, Y.H., Chen, H.I., Wang, Y.Y., Chiou, Y.W., Lin, H.H., Pan, H.A., Wu, C.M., Su, S.M., Hsu, C.C., and Kuo, P.L. (2012). SEPT12 mutations cause male infertility with defective sperm annulus. Hum Mutat 33, 710–719. 10.1002/humu.22028.

59. Lin, Y.H., Wang, Y.Y., Chen, H.I., Kuo, Y.C., Chiou, Y.W., Lin, H.H., Wu, C.M., Hsu, C.C., Chiang, H.S., and Kuo, P.L. (2012). SEPTIN12 genetic variants confer susceptibility to teratozoospermia. PLoS One 7, e34011. 10.1371/journal.pone.0034011.

60. Cavini, I.A., Leonardo, D.A., Rosa, H.V.D., Castro, D., D’Muniz Pereira, H., Valadares, N.F., Araujo, A.P.U., and Garratt, R.C. (2021). The Structural Biology of Septins and Their Filaments: An Update. Front Cell Dev Biol 9, 765085. 10.3389/fcell.2021.765085.

61. Kuo, Y.C., Shen, Y.R., Chen, H.I., Lin, Y.H., Wang, Y.Y., Chen, Y.R., Wang, C.Y., and Kuo, P.L. (2015). SEPT12 orchestrates the formation of mammalian sperm annulus by organizing core octameric complexes with other SEPT proteins. J Cell Sci 128, 923–934. 10.1242/jcs.158998.

62. Mahajan, R., Gerace, L., and Melchior, F. (1998). Molecular characterization of the SUMO-1 modification of RanGAP1 and its role in nuclear envelope association. J Cell Biol 140, 259–270. 10.1083/jcb.140.2.259.

63. Guglielmi, V., Sakuma, S., and D’Angelo, M.A. (2020). Nuclear pore complexes in development and tissue homeostasis. Development 147. 10.1242/dev.183442.

64. Castillo, J., Bogle, O.A., Jodar, M., Torabi, F., Delgado-Duenas, D., Estanyol, J.M., Ballesca, J.L., Miller, D., and Oliva, R. (2019). Proteomic Changes in Human Sperm During Sequential in vitro Capacitation and Acrosome Reaction. Front Cell Dev Biol 7, 295. 10.3389/fcell.2019.00295.

65. Maeshima, K., Iino, H., Hihara, S., Funakoshi, T., Watanabe, A., Nishimura, M., Nakatomi, R., Yahata, K., Imamoto, F., Hashikawa, T., et al. (2010). Nuclear pore formation but not nuclear growth is governed by cyclin-dependent kinases (Cdks) during interphase. Nat Struct Mol Biol 17, 1065–1071. 10.1038/nsmb.1878.

66. Fawcett, D.W., and Chemes, H.E. (1979). Changes in distribution of nuclear pores during differentiation of the male germ cells. Tissue Cell 11, 147–162. 10.1016/0040-8166(79)90015-6.

67. Hayashi, D., Tanabe, K., Katsube, H., and Inoue, Y.H. (2016). B-type nuclear lamin and the nuclear pore complex Nup107-160 influences maintenance of the spindle envelope required for cytokinesis in Drosophila male meiosis. Biol Open 5, 1011–1021. 10.1242/bio.017566.

68. Mostowy, S., and Cossart, P. (2012). Septins: the fourth component of the cytoskeleton. Nature Reviews Molecular Cell Biology 13, 183–194. 10.1038/nrm3284.

69. Mostowy, S., Janel, S., Forestier, C., Roduit, C., Kasas, S., Pizarro-Cerda, J., Cossart, P., and Lafont, F. (2011). A role for septins in the interaction between the Listeria monocytogenes INVASION PROTEIN InlB and the Met receptor. Biophys J 100, 1949–1959. 10.1016/j.bpj.2011.02.040.

70. Barral, Y., Mermall, V., Mooseker, M.S., and Snyder, M. (2000). Compartmentalization of the cell cortex by septins is required for maintenance of cell polarity in yeast. Mol Cell 5, 841–851. 10.1016/s1097-2765(00)80324-x.

71. Szuba, A., Bano, F., Castro-Linares, G., Iv, F., Mavrakis, M., Richter, R.P., Bertin, A., and Koenderink, G.H. (2021). Membrane binding controls ordered self-assembly of animal septins. Elife 10. 10.7554/eLife.63349.

72. Yeh, C.H., Wang, Y.Y., Wee, S.K., Chen, M.F., Chiang, H.S., Kuo, P.L., and Lin, Y.H. (2019). Testis-Specific SEPT12 Expression Affects SUN Protein Localization and is Involved in Mammalian Spermiogenesis. Int J Mol Sci 20. 10.3390/ijms20051163.

73. Andreu, I., Granero-Moya, I., Chahare, N.R., Clein, K., Molina-Jordan, M., Beedle, A.E.M., Elosegui-Artola, A., Abenza, J.F., Rossetti, L., Trepat, X., et al. (2022). Mechanical force application to the nucleus regulates nucleocytoplasmic transport. Nat Cell Biol 24, 896–905. 10.1038/s41556-022-00927-7.

74. Forler, D., Rabut, G., Ciccarelli, F.D., Herold, A., Kocher, T., Niggeweg, R., Bork, P., Ellenberg, J., and Izaurralde, E. (2004). RanBP2/Nup358 provides a major binding site for NXF1-p15 dimers at the nuclear pore complex and functions in nuclear mRNA export. Mol Cell Biol 24, 1155–1167. 10.1128/MCB.24.3.1155-1167.2004.

75. Sabri, N., Roth, P., Xylourgidis, N., Sadeghifar, F., Adler, J., and Samakovlis, C. (2007). Distinct functions of the Drosophila Nup153 and Nup214 FG domains in nuclear protein transport. J Cell Biol 178, 557–565. 10.1083/jcb.200612135.

76. Gorlich, D., and Kutay, U. (1999). Transport between the cell nucleus and the cytoplasm. Annu Rev Cell Dev Biol 15, 607–660. 10.1146/annurev.cellbio.15.1.607.

77. Kierszenbaum, A.L., Gil, M., Rivkin, E., and Tres, L.L. (2002). Ran, a GTP-binding protein involved in nucleocytoplasmic transport and microtubule nucleation, relocates from the manchette to the centrosome region during rat spermiogenesis. Mol Reprod Dev 63, 131–140. 10.1002/mrd.10164.

78. Bayliss, R., Littlewood, T., and Stewart, M. (2000). Structural basis for the interaction between FxFG nucleoporin repeats and importin-beta in nuclear trafficking. Cell 102, 99–108. 10.1016/s0092-8674(00)00014-3.

79. Ori, A., Banterle, N., Iskar, M., Andres-Pons, A., Escher, C., Khanh Bui, H., Sparks, L., Solis-Mezarino, V., Rinner, O., Bork, P., et al. (2013). Cell type-specific nuclear pores: a case in point for context-dependent stoichiometry of molecular machines. Mol Syst Biol 9, 648. 10.1038/msb.2013.4.

80. Raices, M., and D’Angelo, M.A. (2012). Nuclear pore complex composition: a new regulator of tissue-specific and developmental functions. Nat Rev Mol Cell Biol 13, 687–699. 10.1038/nrm3461.

## ADDITIONAL REFERENCES

81. Wagner, F.R., Watanabe, R., Schampers, R., Singh, D., Persoon, H., Schaffer, M., Fruhstorfer, P., Plitzko, J., and Villa, E. (2020). Preparing samples from whole cells using focused-ion-beam milling for cryo-electron tomography. Nat Protoc 15, 2041–2070. 10.1038/s41596-020-0320-x.

82. Sexton, D.L., Burgold, S., Schertel, A., and Tocheva, E.I. (2022). Super-resolution confocal cryo-CLEM with cryo-FIB milling for in situ imaging of Deinococcus radiodurans. Curr Res Struct Biol 4, 1–9. 10.1016/j.crstbi.2021.12.001.

83. Eisenstein, F., Yanagisawa, H., Kashihara, H., Kikkawa, M., Tsukita, S., and Danev, R. (2023). Parallel cryo electron tomography on in situ lamellae. Nat Methods 20, 131–138. 10.1038/s41592-022-01690-1.

84. Hagen, W.J.H., Wan, W., and Briggs, J.A.G. (2017). Implementation of a cryo-electron tomography tilt-scheme optimized for high resolution subtomogram averaging. J Struct Biol 197, 191–198. 10.1016/j.jsb.2016.06.007.

85. Tegunov, D., and Cramer, P. (2019). Real-time cryo-electron microscopy data preprocessing with Warp. Nat Methods 16, 1146–1152. 10.1038/s41592-019-0580-y.

86. Zheng, S., Wolff, G., Greenan, G., Chen, Z., Faas, F.G.A., Barcena, M., Koster, A.J., Cheng, Y., and Agard, D.A. (2022). AreTomo: An integrated software package for automated marker-free, motion-corrected cryo-electron tomographic alignment and reconstruction. J Struct Biol X 6, 100068. 10.1016/j.yjsbx.2022.100068.

87. Kremer, J.R., Mastronarde, D.N., and McIntosh, J.R. (1996). Computer visualization of three-dimensional image data using IMOD. J Struct Biol 116, 71–76. 10.1006/jsbi.1996.0013.

88. Burt, A. (2024). Script for converting dipole annotations as IMOD model files into oriented particles compatible with Warp. Zenodo. 10.5281/zenodo.14010295.

89. Zivanov, J., Nakane, T., Forsberg, B.O., Kimanius, D., Hagen, W.J., Lindahl, E., and Scheres, S.H. (2018). New tools for automated high-resolution cryo-EM structure determination in RELION-3. Elife 7. 10.7554/eLife.42166.

90. Dendooven, T., Yatskevich, S., Burt, A., Chen, Z.A., Bellini, D., Rappsilber, J., Kilmartin, J.V., and Barford, D. (2024). Structure of the native gamma-tubulin ring complex capping spindle microtubules. Nat Struct Mol Biol 31, 1134–1144. 10.1038/s41594-024-01281-y.

91. Castano-Diez, D., Kudryashev, M., Arheit, M., and Stahlberg, H. (2012). Dynamo: a flexible, user-friendly development tool for subtomogram averaging of cryo-EM data in high-performance computing environments. J Struct Biol 178, 139–151. 10.1016/j.jsb.2011.12.017.

92. Burt, A., Gaifas, L., Dendooven, T., and Gutsche, I. (2021). A flexible framework for multi-particle refinement in cryo-electron tomography. PLoS Biol 19, e3001319. 10.1371/journal.pbio.3001319.

93. Tegunov, D., Xue, L., Dienemann, C., Cramer, P., and Mahamid, J. (2021). Multi-particle cryo-EM refinement with M visualizes ribosome-antibiotic complex at 3.5 A in cells. Nat Methods 18, 186–193. 10.1038/s41592-020-01054-7.

94. Campello, R.J.G.B., Moulavi, D., and Sander, J. (2013). Density-Based Clustering Based on Hierarchical Density Estimates. held in Berlin, Heidelberg, 2013//. J. Pei, V.S. Tseng, L. Cao, H. Motoda, and G. Xu, eds. (Springer Berlin Heidelberg), pp. 160-172.

